# Cognitive reserve depends on cortical neurochemistry to predict memory after partner-inflicted brain injury

**DOI:** 10.64898/2026.10.27.762370

**Authors:** Craig W. McFarland, Alexander P. Lin, Skyler McComas, Eva-Maria Ratai, Atilla Gönenç, Annie-Lori Joseph Denk, Sofia Nikolaidou, Eve M. Valera

## Abstract

Memory problems are commonplace following brain injury (BI). However, it remains unclear why some report memory problems while others do not, especially following repeated neurotrauma. Cognitive reserve has been widely hypothesized to account for this variation, but testing this account has yielded inconsistent results and proven difficult, likely as it requires a sample spanning a wide range of BI burden with concurrent neuroimaging and cognitive measurement. Here, we combined a wide range of cognitive assessments with measures of neuronal health using magnetic resonance spectroscopy across three brain regions in 62 women who had experienced physical intimate partner violence. Partner-inflicted BI counts ranged from 0 to 420, yet we found that the number of BIs alone did not explain the variance in memory outcomes. Among a small subset with high BI exposure (≥10 BIs; *n* = 11), BI count explained 5% of the variance in delayed recall, the primary memory outcome, whereas cognitive reserve, inferred from a measure of premorbid functioning, explained almost half (unadjusted in-sample associations). In the full cohort, cognitive reserve predicted memory only among the most heavily injured participants. In a separate model, cognitive reserve predicted memory only when creatine-referenced total N-acetylaspartate, a marker of neuronal health, was lower in the posterior cingulate gyrus and dorsolateral prefrontal cortex. Ultimately, our findings provide behavioral and neurochemical evidence consistent with the cognitive reserve hypothesis and shed new light on why, despite a high burden of neurotrauma, memory may remain resilient.

## Introduction

Memory problems are commonplace among those who sustain a brain injury (BI) [1–3]. This is also evident in the case of physical intimate partner violence (IPV), where survivors often experience blows to the head and strangulations that can injure the brain [4–8], which in turn may impair memory. Relatedly, IPV and partner-inflicted BI have been associated with poorer performance on tests of memory, among other cognitive domains [9–14], as well as with brain structure and connectivity in memory-related circuitry [15–18]. Moreover, survivors of IPV often sustain multiple BIs over years of abuse, and these “invisible” injuries rarely reach medical attention [19,20].

Cognitive outcomes after BI are not uniform. For instance, some individuals who report multiple partner-inflicted BIs, including BIs involving strangulation or loss of consciousness, perform at or above population norms, while others present learning and memory deficits [9,10]. Why memory appears to remain resilient in some cases and yet become demonstrably impaired in others remains unclear.

Cognitive reserve, defined as the adaptability of cognitive processes to maintain performance despite brain damage or pathology, has been widely hypothesized to account for vastly different cognitive outcomes [21–23]. For instance, in older adults with Alzheimer’s disease, cognitive reserve modified the association between Alzheimer’s disease pathology and cognitive function, such that those with more education performed better at comparable levels of pathology [24]. Notably, cognitive reserve differs from brain reserve, which reflects the brain’s neurobiological capacity rather than the adaptability of the cognitive processes it supports [25]. Cognitive reserve is commonly estimated from education, occupational complexity, leisure, and premorbid functioning [22], and modifies the association between brain status (i.e., the extent of brain change, injury, or pathology) and cognitive performance as opposed to merely covarying with cognitive performance [22,25]. This idea was first suggested by earlier neuropathological evidence of preserved cognition despite substantial pathology [26].

Testing these predictions in the context of BI has proven difficult because the ideal design requires a cohort spanning a wide range of neurotrauma burden alongside concurrent measures of cognitive reserve, brain status, and cognition. It therefore remains unclear under which conditions cognitive reserve protects memory following BI and which neurobiological conditions influence that protection. Examining memory among survivors of repeated partner-inflicted BI, whose injury burden varies widely, can help identify the neurochemical conditions under which cognitive reserve protects memory.

Here, we combined cognitive assessments with neuroimaging in 62 women with a history of physical IPV, whose partner-inflicted BIs ranged from 0 to 420 (see Methods). Cumulative partner-inflicted BI was characterized by structured interviews. Cognitive assessment included examiner-administered neuropsychological testing and a computerized battery of ten cognitive tests yielding fifteen domain scores. In our neuroimaging sample, proton magnetic resonance spectroscopy (MRS) provided in vivo measures of brain neurochemistry to examine the biological context of memory preservation. We focused primarily on the ratio of total N-acetylaspartate to total creatine (tNAA/tCr), which is reliably quantified and frequently reduced after TBI [27], in three regions: the posterior cingulate gyrus (PCG), which shows injury-related NAA reductions [28] and connectivity associated with learning and memory after partner-inflicted BI [20]; the dorsolateral prefrontal cortex (DLPFC), which contributes to working memory [29]; and the corona radiata, a white-matter region in which lesions have been associated with verbal memory impairment [30]. The remaining metabolites were examined as exploratory analyses.

Ultimately, we aimed to determine whether cognitive reserve, operationalized as performance on a measure of estimated premorbid functioning, predicts memory following BI and, if so, under what neurochemical conditions. In our cohort of women with a history of physical IPV, identifying these conditions may help clarify the neurobiology of memory preservation following repeated partner-inflicted BI. Understanding cognitive reserve may help reveal the biological mechanisms of memory resilience and inform future interventions to preserve or restore memory.

## Results

### Brain injury burden does not explain the majority of memory outcomes

We first characterized BI burden in the cohort (Table 1). Over two thirds (68%, *n* = 42) had sustained at least one partner-inflicted BI, inclusive of BI due to blunt force, strangulation, and violent shaking, with a median of 3 among those with any BI (interquartile range 1 to 10) and a maximum of 420 (see Methods). Eighteen participants (29%) reported five or more BIs and twelve (19%) reported ten or more.

**Table 1.** Participant characteristics by number of partner-inflicted brain injuries.

| **Characteristic** | All (*n* = 62) | **Partner-inflicted brain injuries** |  |  | *P* |
|---|---|---|---|---|---|
|  |  | 0 (*n* = 20) | 1–9 (*n* = 30) | ≥10 (*n* = 12) |  |
| Age, years | 40.2 (13.1) | 37.8 (13.8) | 42.1 (12.7) | 39.5 (13.1) | 0.472 |
| Education, years | 16.5 (2.6) | 17.9 (2.3) | 16.0 (2.6) | 15.5 (2.2) | 0.015 |
| Race other than White, *n* (%) | 17 (27) | 4 (20) | 11 (37) | 2 (17) | 0.287 |
| Hispanic or Latina, *n* (%) | 10 (17) | 2 (11) | 6 (21) | 2 (17) | 0.657 |
| Cognitive reserve (ToPF standard score) | 108.9 (10.8),<br>76–124 | 109.0 (8.7),<br>86–118 | 107.8 (13.2),<br>76–124 | 111.5 (7.3),<br>98–120 | 0.763 |
| Partner-inflicted brain injuries, median (IQR) | 1 (0–5) | 0 (0–0) | 2 (1–4) | 15 (13–27) | <.001 |
| Years since last partner-inflicted injury | 11.3 (8.8) | — | 12.7 (9.3) | 7.9 (6.8) | 0.122 |
| Post-traumatic stress (PCL-5) | 30.4 (17.6) | 30.9 (20.6) | 27.3 (15.8) | 37.2 (15.8) | 0.154 |
| Depression (PHQ-9) | 8.3 (5.3) | 8.3 (5.8) | 7.7 (4.6) | 9.8 (6.3) | 0.702 |
| Alcohol use (AUDIT) | 3.1 (4.3) | 2.7 (3.7) | 3.2 (4.0) | 3.6 (5.9) | 0.759 |
| Childhood maltreatment (CTQ) | 53.8 (21.2) | 48.0 (18.3) | 56.8 (25.0) | 55.8 (13.0) | 0.291 |
| Abuse severity, past year (CAS) | 7.3 (11.9) | 11.6 (15.4) | 3.7 (6.1) | 9.0 (14.5) | 0.284 |

Values are mean (SD) unless otherwise stated, and premorbid functioning is given as mean (SD), range. Cognitive reserve was estimated with the Test of Premorbid Functioning (ToPF). *P* values are Kruskal-Wallis tests across the three exposure groups. Years since last partner-inflicted injury is accordingly undefined for women reporting no injury. Twelve participants reported ten or more injuries, of whom eleven completed delayed recall. Sample sizes otherwise vary because not every measure was available for every participant, and the analytic *n* for each model is given in Supplementary Table 1.

Next, we characterized memory outcomes. Across the full cohort, performance on the examiner-administered (CVLT-3) and computerized (CNS Vital Signs) memory tests ranged from the 2nd to the 98th percentile of the general population. Among participants with the greatest burden of neurotrauma (≥10 BIs), in whom we expected the most memory impairment, delayed-recall performance (i.e., the ability to recall previously learned information after a delay) ranged from the 2nd to the 91st percentile, with a spread (SD 3.27) similar to that of the general population (SD 3.00).

Memory outcomes diverged even at identical reported injury counts. To illustrate, two participants who each reported exactly ten BIs scored at the 25th and 75th percentiles on the same delayed recall task. This heterogeneity in memory performance appeared on every memory measure administered, across both the examiner-administered and the computerized battery (Fig. 1 and Supplementary Table 1).

**Fig. 1.**
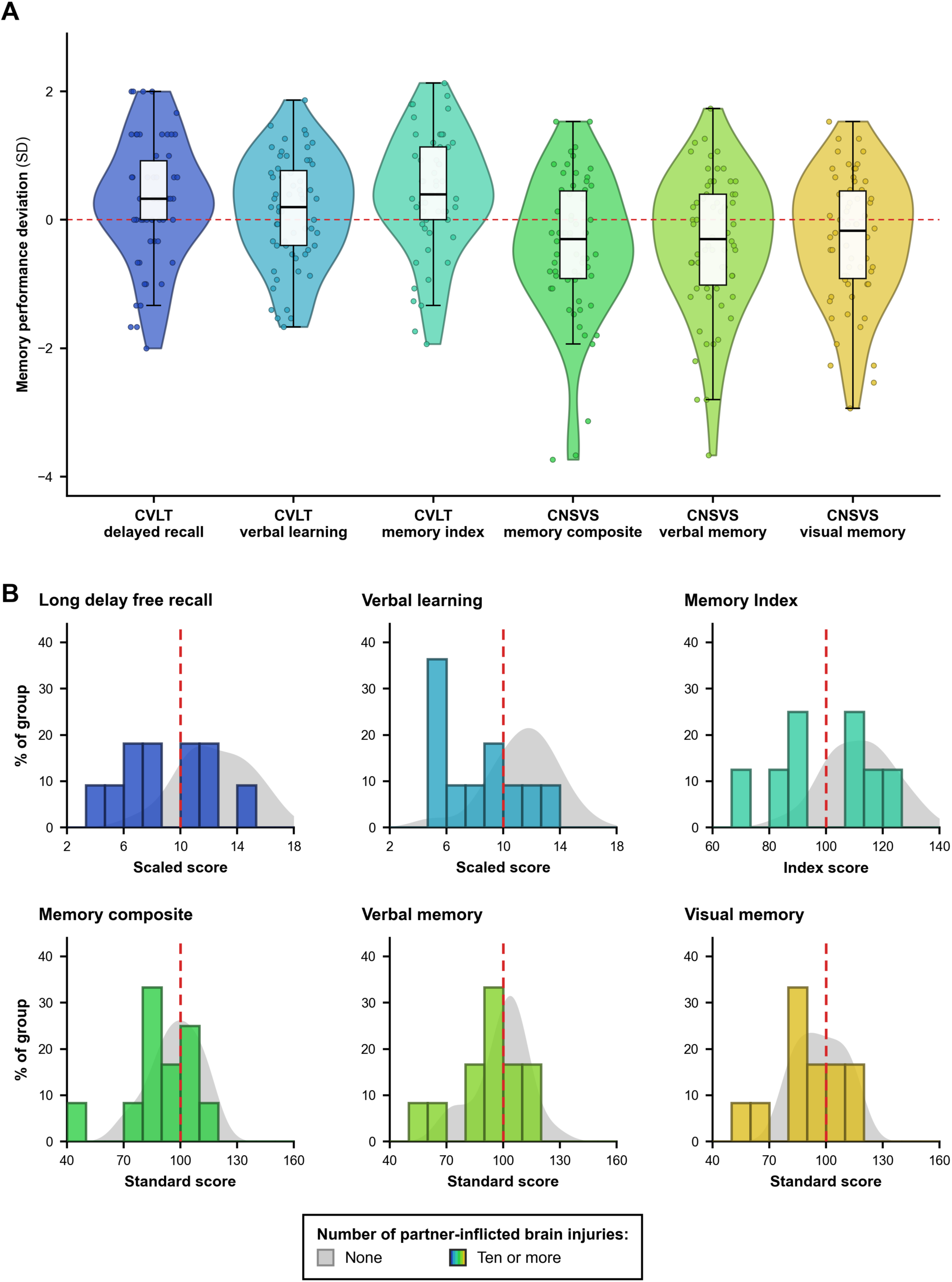
Memory outcomes vary widely across injury burden, including among participants with the greatest burden of partner-inflicted neurotrauma. (A) Distributions of six memory measures across the full cohort, standardized relative to published normative data. Zero denotes the normative mean and one unit represents one normative standard deviation. Violin plots show density, boxes show medians and interquartile ranges, and points represent individual participants. (B) Distributions of scores on each measure’s original scale for participants reporting no partner-inflicted BI and those reporting ten or more injuries, each panel headed with the measure name and presented in the same order as in (A). Colored histograms represent the group reporting ten or more injuries, and gray density curves represent the group reporting no BI. Distributions are normalized within exposure group to account for unequal group sizes. Red dashed lines mark the published normative means. The groups overlapped substantially across every measure, with memory performance in the highest-burden group extending from impaired levels to above the normative mean. CVLT-3, California Verbal Learning Test 3rd Edition; CNSVS, CNS Vital Signs.

Then, we asked whether the number of BIs reported could account for this variation in memory outcomes. Across the full cohort, participants who reported more Bis demonstrated significantly poorer long-delay free recall (*β* = −0.49, 95% CI [−0.75, −0.23], *P* = .0004), adjusting for education. Yet, BI burden accounted for at most 27.0% of the variance in any memory measure (unadjusted Pearson r2; Supplementary Table 2). Moreover, among participants with the greatest burden of neurotrauma (≥10 BIs), the number of BIs did not account for memory outcomes. Squared rank correlations between BI burden and memory ranged from only 0.2% to 12.2% across measures (unadjusted Pearson *r^2^*, 0.1% to 25.1%), and no association was detected on any measure (all *P* ≥ .29; Supplementary Table 2). Thus, although more BIs were associated with poorer memory overall, the number of BIs did not explain why memory varied so widely among those with the greatest burden of neurotrauma.

### Cognitive reserve predicts memory only at the highest injury burden

Next, we hypothesized that cognitive reserve may explain the memory resilience observed within the cohort. Specifically, participants with greater cognitive reserve should show better memory performance, irrespective of how many injuries a participant had sustained. Contrary to expectation, cognitive reserve was not significantly associated with delayed recall, the primary outcome, after adjustment (*β* = 0.10 to 0.11, all *P* ≥ .43), or with any other memory measure. The associations closest to conventional significance were with working memory (*β* = 0.27, 95% CI [−0.003, 0.55], *P* = .053) and visual memory (*β* = 0.29, 95% CI [−0.03, 0.61], *P* = .078); all other P values were ≥ .16.

Threshold accounts of brain reserve hold that cognitive impairment emerges only once injury or pathology exceeds what neurobiological capacity can absorb [31]. To test this hypothesis, we examined the same association among participants with the greatest neurotrauma exposure (≥10 BIs), and found that greater cognitive reserve indeed strongly predicted memory in these participants (Spearman *ρ* = 0.73, *P* = .011, *n* = 11). Given the small size of this group, we performed a leave-one-out analysis, and the association remained significant and similar in magnitude (*ρ* = 0.63 to 0.86, all *P* ≤ .049). Notably, the positive association between cognitive reserve and delayed recall was specific to those who had sustained the most BIs, as cognitive reserve was not significantly associated with delayed recall among participants reporting one to nine BIs (*ρ* = 0.14, *P* = .46, *n* = 28) or no partner-inflicted BI (*ρ* = −0.23, *P* = .34, *n* = 19; Fig. 2A, 2B). Beyond delayed recall, the reserve-by-injury interaction was also present for CNSVS visual memory (*<u>β</u>* = 1.38, 95% CI [0.64, 2.12], *P* = .0004) and the CNSVS memory composite (*β* = 1.30, 95% CI [0.03, 2.56], *P* = .045), but not for CNSVS verbal or working memory (*P* = .31 and .47, respectively).

**Fig. 2.**
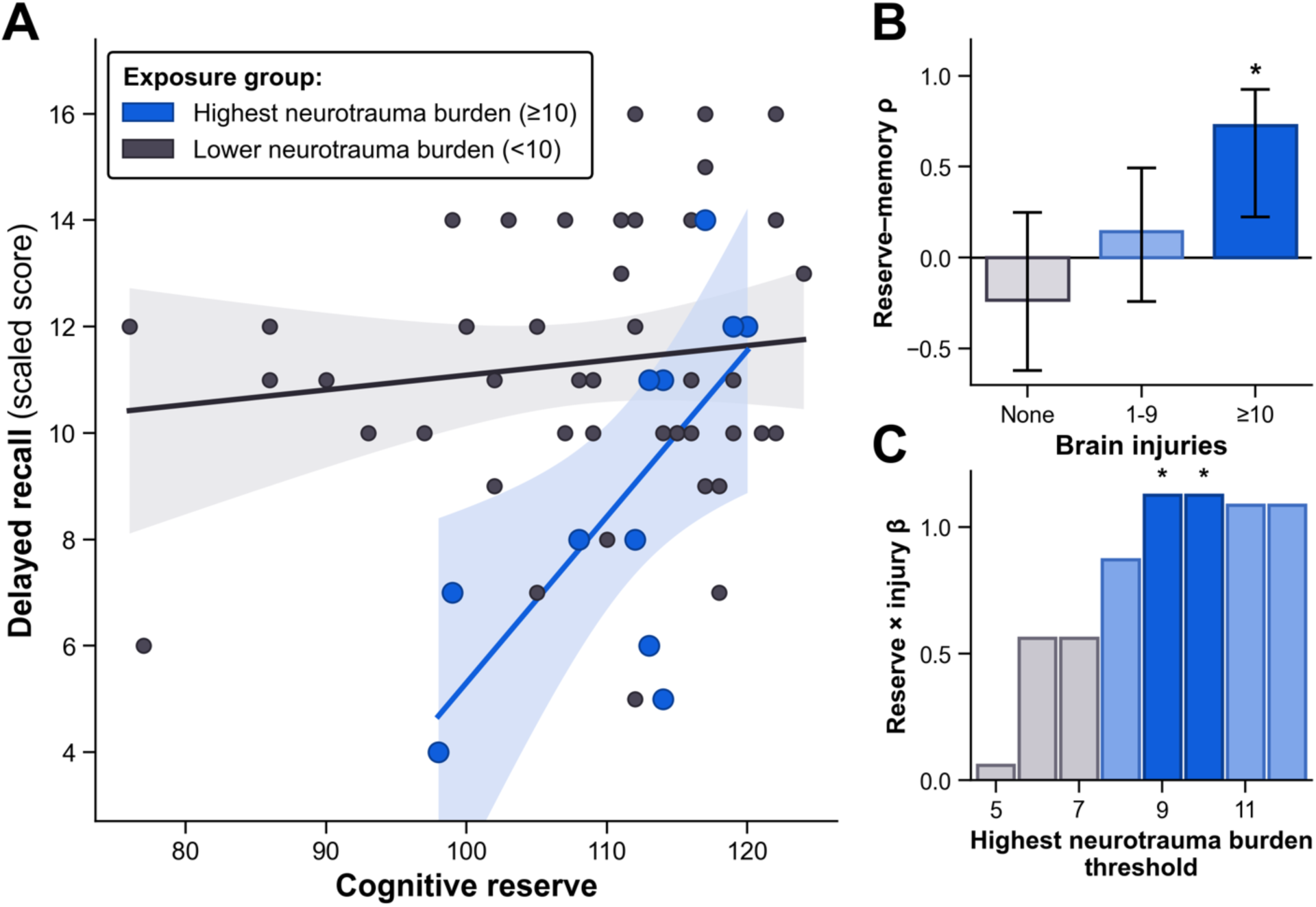
Cognitive reserve predicts memory only among participants with the greatest injury burden. **(A)** Cognitive reserve, measured by the Test of Premorbid Functioning standard score, plotted against California Verbal Learning Test-3 long-delay free recall for participants reporting ten or more partner-inflicted brain injuries (blue) and fewer than ten partner-inflicted brain injuries (gray). Lines show separate least-squares fits and shaded regions show 95% confidence bands. Spearman’s ρ = 0.73 (*P* = .011; n = 11) in the highest-burden group and ρ = 0.02 (*P* = .91; n = 47) among participants reporting fewer than ten injuries. Greater cognitive reserve was associated with better memory only in the highest-burden group. **(B)** Spearman correlations between cognitive reserve and delayed recall among participants reporting no injuries, one to nine injuries and ten or more injuries. Error bars show 95% confidence intervals derived using Fisher z-transformation. The association was detected only in the highest exposure stratum. **(C)** Reserve-by-injury interaction coefficients from the binary model (highest band against all others; Supplementary Table 4) across candidate thresholds of five to twelve injuries. Blue bars indicate *P* < .05, paler blue bars .05 ≤ *P* < .10, and gray bars *P* ≥ .10. The interaction estimate rises from 0.06 at five injuries to 1.13 at nine and ten injuries and is similar (1.09) at eleven and twelve, where the highest-burden band contains only nine participants. No participant reported exactly six, nine or eleven injuries, resulting in identical groups at the corresponding adjacent thresholds. In the full cohort without stratification by injury burden, no association between cognitive reserve and delayed recall was detected. \**P* < .05.

Moreover, among those with the greatest neurotrauma exposure, a one SD advantage in cognitive reserve corresponded to 1.25 SD of better delayed recall performance (*β* = 1.25, 95% CI [0.42, 2.09], *P* = .004). This advantage corresponds approximately to the difference between placing at the 11th and 50th percentiles of the general population. Notably, the association between cognitive reserve and delayed recall was significantly stronger among women with ten or more BIs than among those with none (*β* = 1.36, 95% CI [0.23, 2.49], *P* = .019), and among participants with no partner-inflicted BI, a one standard deviation advantage in cognitive reserve was not associated with a significant gain in delayed recall (*β* = −0.11, 95% CI [−0.81, 0.59], *P* = .75).

To test the robustness of this finding, we assessed whether the reserve-by-injury interaction depended on individual participants, on the cutoff used to define the highest-burden group, or on the covariates included. In a leave-one-out analysis, the interaction remained similar in magnitude (*β* = 1.16 to 1.67), although it was no longer significant when any one of four participants was omitted (maximum *P* = .17). A Freedman–Lane permutation test, which does not rely on distributional assumptions, supported the interaction (*P* = .024; Supplementary Table 3), and adding the interaction improved out-of-sample prediction of delayed recall in leave-one-out cross-validation (cross-validated *R^2^* from 0.01 to 0.07). In the primary three-strata model, the interaction held at every threshold between nine and twelve BIs. These four thresholds resolve to only two distinct partitions of the cohort, placing either eleven participants (*β* = 1.36, 95% CI [0.23, 2.49], *P* = .019) or nine participants (*β* = 1.32, 95% CI [0.06, 2.57], *P* = .040) in the highest band; when the highest band was instead contrasted with all other participants, the corresponding estimates were *β* = 1.13 (*P* = .021) and *β* = 1.09 (*P* = .056; Fig. 2C and Supplementary Table 4). The interaction was not observed when injury burden was coded continuously (Fig. S1). The interaction also held after additional adjustment for depression symptoms, childhood maltreatment, alcohol use [32], other sources of BI and past-year abuse severity on the Composite Abuse Scale (*β* = 1.23 to 1.40, all *P* ≤ .032; Supplementary Table 3). No difference in the interaction was detected between injury mechanisms (blunt force versus strangulation, difference *β* = 0.26, 95% CI [−0.27, 0.80], *P* = .33; Supplementary Table 5).

Finally, to determine whether the reserve-by-injury interaction was specific to memory as opposed to cognitive performance more generally, we repeated the analysis across non-memory domains of the computerized battery (e.g., attention, processing speed, executive function and reasoning). The interaction was not present for processing speed (*β* = −0.19, 95% CI [−1.00, 0.62], *P* = .64) or executive function (*β* = −0.20, 95% CI [−1.90, 1.50], *P* = .82), two domains commonly affected after BI [2]. Across all fifteen domains, the interaction was significantly stronger in memory than in non-memory domains (*β* = 1.12, 95% CI [0.68, 1.57], *P* < .001; Supplementary Table 6). When education, the most widely used alternative proxy for cognitive reserve, was substituted for premorbid functioning, its interaction with injury burden was not significant (*β* = 0.33, 95% CI [−1.04, 1.70], *P* = .63).

### Cortical tNAA/tCr moderates the association between cognitive reserve and memory

Brain reserve, the brain’s neuroanatomical resources that afford greater capacity for cognitive resilience, has been proposed to serve as a structural capacity to maintain cognitive function in spite of injury [25]. Total N-acetylaspartate (tNAA) to total creatine (tCr) may be one proxy for brain reserve, as tNAA/tCr serves as a marker of neuronal and metabolic health [33]. We therefore examined whether tNAA/tCr moderates the association between cognitive reserve and memory. Indeed, tNAA/tCr in the PCG moderated the relationship between cognitive reserve and delayed recall (*β* = −0.46, 95% CI [−0.71, −0.20], *P* = .0008) after adjustment (Fig. 3C). The effect survived Benjamini-Hochberg correction across the three brain regions in which tNAA/tCr was measured (*q* = .002).

**Fig. 3.**
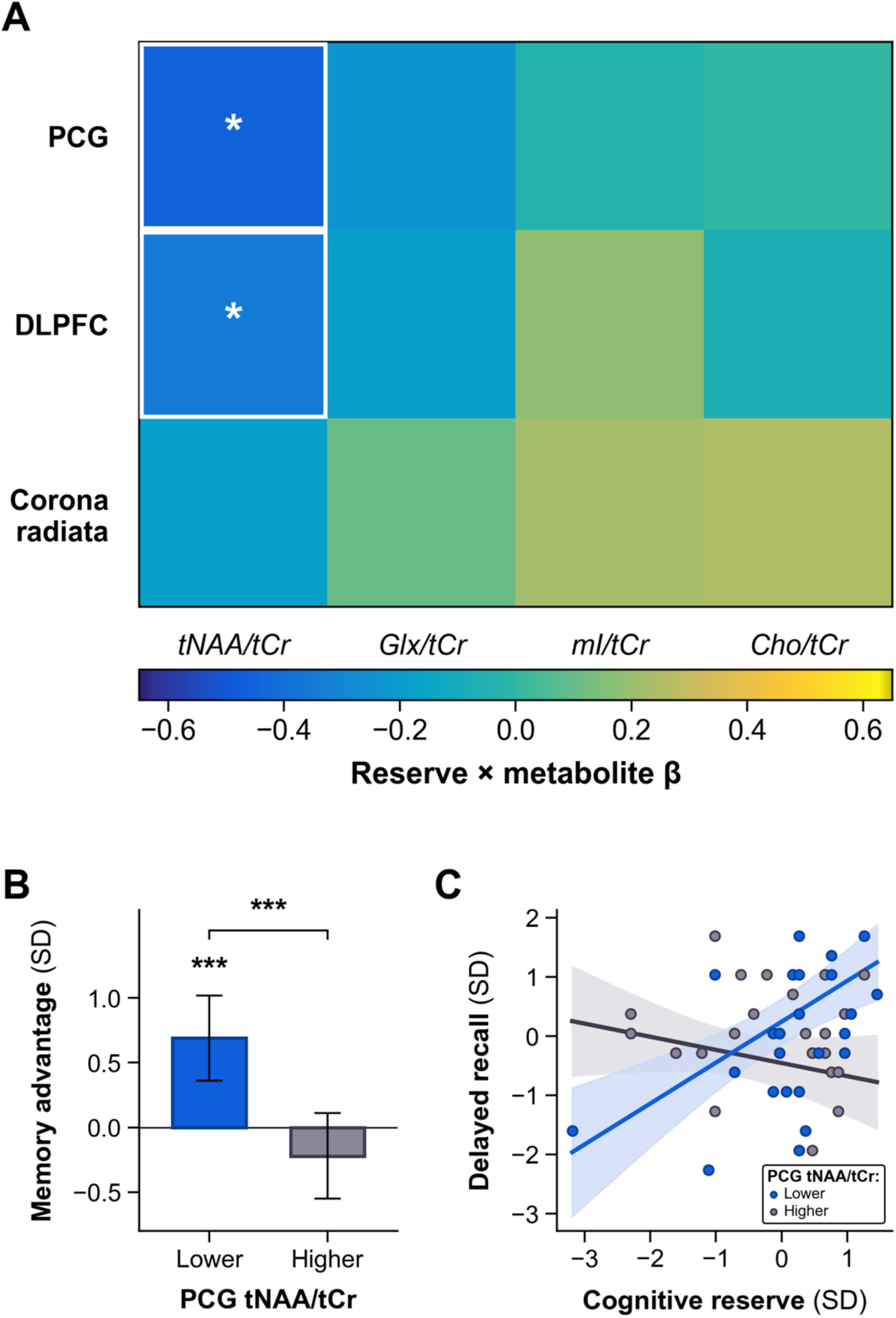
Cognitive reserve predicts memory only at lower PCG tNAA/tCr. **(A)** Heatmap of cognitive reserve-by-metabolite interaction coefficients for the twelve metabolite and voxel combinations examined. Rows represent MRS voxels and columns represent metabolites, with all metabolite concentrations expressed relative to total creatine. The color scale denotes the interaction coefficient, with negative values indicating stronger positive associations between cognitive reserve and memory at lower metabolite concentrations. Cells outlined in white and marked with an asterisk survived Benjamini-Hochberg false-discovery-rate correction and confidence intervals are reported in Supplementary Table 7. Six spectra were excluded at quality control, two from the DLPFC voxel and four from the corona radiata, and four participants lacked delayed recall, so the adjusted models include 47 participants for the PCG, 46 for the DLPFC and 45 for the corona radiata. **(B)** Estimated difference in delayed verbal recall associated with a one-standard-deviation higher cognitive reserve score at lower PCG tNAA/tCr (1 SD below the mean, 1.29) and higher PCG tNAA/tCr (1 SD above the mean, 1.48). Error bars show 95% confidence intervals. Estimates above zero indicate better recall at higher cognitive reserve. An association was detected at lower tNAA/tCr but not at higher tNAA/tCr. **(C)** Cognitive reserve plotted against delayed recall. Observations are grouped at the median of PCG tNAA/tCr (1.37; *n* = 24 lower, 23 higher) for visualization. Lines show predictions from a single model treating tNAA/tCr as continuous, evaluated at one standard deviation below (1.29) and above (1.48) its mean. Shaded regions show 95% confidence bands. All inference used the continuous moderator. PCG, posterior cingulate gyrus; DLPFC, dorsolateral prefrontal cortex; Glx, glutamate plus glutamine; mI, myo-inositol; Cho, choline-containing compounds; tNAA/tCr, total N-acetylaspartate-to-total-creatine ratio. \**P* < .05, \*\**P* < .01, \*\*\**P* < .001.

In addition, cognitive reserve predicted delayed recall (*β* = 0.69, 95% CI [0.36, 1.02], *P* = .0001) among participants who had a PCG tNAA/tCr ratio 1 SD below the sample mean, but not among those 1 SD above (*β* = −0.22, 95% CI [−0.55, 0.11], *P* = .18; Fig. 3A and B). PCG tNAA/tCr averaged 1.39 (SD 0.10; range 1.22–1.64), so these conditioning values correspond to ratios of 1.29 and 1.48. tNAA/tCr was approximately normally distributed in all three voxels (Shapiro–Wilk *P* ≥ .15; Supplementary Table 7).

The interaction was robust to bootstrap resampling, to the omission of any single participant and to the inclusion of additional covariates, and adding it improved out-of-sample prediction of delayed recall (cross-validated R² from 0.09 to 0.33; Supplementary Table 8). The interaction was also disordinal: tNAA/tCr was not associated with recall at lower cognitive reserve (1 SD below the mean; *β* = 0.11, 95% CI [−0.23, 0.45], *P* = .53) but was inversely associated with recall at higher cognitive reserve (1 SD above the mean; *β* = −0.80, 95% CI [−1.27, −0.33], *P* = .001; Fig. S2).

To determine whether tNAA/tCr and injury burden each moderated the association between cognitive reserve and memory when considered together, we entered both moderators in a single model of the same 47 participants. Both moderators remained significant (PCG tNAA/tCr *β* = −0.42, 95% CI [−0.69, −0.15], *P* = .003; ten or more partner-inflicted BIs, entered as a binary indicator, *β* = 0.70, 95% CI [0.01, 1.39], *P* = .046), with the metabolite ratio the more precisely estimated of the two. Two further features distinguish the metabolite from a marker of accumulated damage. The metabolite ratio was not significantly associated with delayed recall (*ρ* = −0.001, *P* = .99) or with the number of partner-inflicted BIs (*ρ* = −0.11, *P* = .43), although these null marginal associations do not exclude relationships with memory or injury burden.

As the moderator is a ratio, we also entered tNAA and tCr separately, each with its own moderation term. Neither component produced the interaction alone (tNAA *β* = −0.27, 95% CI [−0.63, 0.09], *P* = .14; tCr *β* = 0.22, 95% CI [−0.21, 0.65], *P* = .32), and with both moderation terms in the model the tNAA term was negative (*β* = −0.45, 95% CI [−0.77, −0.14], *P* = .006) and the tCr term positive (*β* = 0.39, 95% CI [0.06, 0.72], *P* = .020). Given that a change in significance after mutual adjustment does not identify which component is biologically primary, the moderation is attributed to the ratio rather than to either component. The three-way interaction between cognitive reserve, PCG tNAA/tCr and the highest-burden band was not significant (*β* = −0.17, 95% CI [−1.51, 1.17], *P* = .80), so this cohort cannot establish that the expression of cognitive reserve depends on high injury burden and lower tNAA/tCr simultaneously.

To establish whether this pattern was confined to one memory measure or one brain region, we repeated the tNAA/tCr interaction using other memory measures and in the two other brain regions. The same interaction was present on a composite of verbal learning and delayed recall (*β* = −0.37, 95% CI [−0.64, −0.11], *P* = .007) and on visual memory (*β* = −0.24, 95% CI [−0.47, −0.01], *P* = .040) of the computerized battery. The same pattern appeared for DLPFC tNAA/tCr, where the interaction (*β* = −0.35, 95% CI [−0.65, −0.05], *P* = .024) survived FDR correction (*q* = .036) and cognitive reserve predicted memory at lower values (*β* = 0.55, 95% CI [0.08, 1.02], *P* = .024), although this interaction was less robust than that in the PCG, with leave-one-out estimates ranging from *β* = −0.40 to −0.26 and *P* values up to .15. Choline in the corona radiata showed the same conditional pattern in the opposite metabolite direction, with cognitive reserve predicting better memory at higher choline (conditional slope *β* = 0.54, 95% CI [0.18, 0.89], *P* = .004; interaction *β* = 0.26, 95% CI [−0.01, 0.53], *P* = .063), although this interaction was not statistically significant. None of the three metabolites was independently associated with memory or with BI burden (all *P* ≥ .33). Myo-inositol, a biomarker of glial activity, showed no significant interaction in any brain region (all *P* ≥ .20).

## Discussion

In this study, we combined a wide range of cognitive assessments with proton MRS in a cohort of 62 women with a history of physical IPV to investigate why, despite a high burden of neurotrauma, memory is impaired in some yet remains resilient in others. To address this question, we tested whether the cognitive reserve hypothesis could explain this variation.

Among participants with the greatest burden of neurotrauma (*n* = 11), BI burden explained 5% of the variance in delayed recall, the primary memory outcome, whereas cognitive reserve explained 48% (unadjusted Pearson *r^2^*). Across all eight memory measures, the corresponding values ranged from 0.1% to 25.1% for BI burden and from 22.7% to 75.0% for cognitive reserve (Supplementary Table 2). Greater cognitive reserve also predicted better memory, but only among a small subset of participants with the greatest cumulative neurotrauma burden (≥10 BIs; n = 11), a finding that warrants replication in larger samples. Independently, higher cognitive reserve predicted better memory only at lower levels of tNAA/tCr, commonly interpreted as an indicator of neuronal and metabolic health, within the PCG. Together, these two conditional effects are consistent with the consensus operational definition of cognitive reserve, in which a reserve proxy moderates the association between brain status and cognition [22]. As such, in our sample, cognitive reserve predicted better memory most strongly in participants who had the highest burden of neurotrauma and, in a separate analysis, in those with relatively lower tNAA/tCr. Taken together, these behavioral and neurochemical findings are consistent with the cognitive reserve hypothesis and point to a neurobiological context in which reserve is expressed.

### Evidence consistent with cognitive reserve after brain injury

To our knowledge, this study provides the first evidence that cognitive reserve, the capacity to preserve cognition despite BI, predicts better memory among women with partner-inflicted BI, depending on injury burden and PCG tNAA/tCr. Previous studies used estimated premorbid functioning either to quantify deviation from expected functioning [10] or as a covariate in cognitive comparisons [9]. Here, cognitive reserve interacted with PCG tNAA/tCr in predicting memory. This finding is directly in line with what consensus criteria identify as the ideal demonstration of cognitive reserve, namely that a cognitive reserve proxy should be shown to moderate the relationship between an indicator of brain status (e.g., tNAA/tCr) and cognition (e.g., memory), so that cognition is predicted by the interaction between the two [22].

Leveraging behavioral and neurochemical measures in a cohort spanning 0 to 420 BIs allowed us to perform this moderation test. MRS offered neurochemical evidence, as higher cognitive reserve predicted better memory only when PCG tNAA/tCr was lower, with lower values commonly interpreted as evidence of poorer neuronal or metabolic health after TBI [27,28]. The consensus framework further notes that direct examination of neural mechanisms, rather than sociocultural proxies alone, may yield more insight into how cognitive reserve operates [25], and pairing a reserve proxy via neuropsychological testing with MRS permitted us to do both.

### A window model of cognitive reserve

Considered alongside emerging neuroimaging evidence, our data suggest that cognitive reserve may be most relevant to memory within a “window” of neurobiological compromise. Our study points to a possible minimum level of neurobiological compromise, at which a positive association between cognitive reserve and delayed recall was detected only among participants with the greatest cumulative BI burden (≥10 BIs; n = 11), suggesting that sufficient neurobiological challenge may be needed before differences in reserve become apparent. Krch et al. [34] point to a possible maximum, as, in their predominantly severe TBI cohort, the memory advantage associated with cognitive reserve was greatest at lower levels of DTI-defined neuropathology, weakened as DTI-defined neuropathology increased, and was no longer detected at the highest levels. Critically, their cohort predominantly included severe TBI, whereas ours excluded moderate or severe TBI, so differences in injury severity prevent direct comparison of the proposed boundaries. Finally, Vockert et al. [35] provided further evidence for a state-dependent effect of cognitive reserve, where brain activity associated with cognitive reserve (e.g., stronger deactivation of core cognitive circuits) weakened the relationship between Alzheimer’s disease pathology and poorer cognition.

Together, these findings suggest that cognitive reserve does not confer a uniform benefit across levels of brain pathology. When neurobiological capacity is sufficient for cognitive demands, reserve may contribute little additional benefit. As that capacity becomes strained, individuals with greater reserve may be better able to maintain memory through adaptive recruitment or more efficient use of remaining resources [21,22]. With extensive damage, however, the capacity for such adaptation may become constrained. Cognitive reserve may therefore operate within a window, contributing most when neurobiological capacity is compromised but remains sufficient for adaptation [36]. A systematic review of 55 imaging-biomarker studies in Alzheimer’s disease similarly reports that the benefit of cognitive reserve depends on disease stage, diminishing and in some studies reversing as pathology advances [37].

Our results therefore extend previous BI studies, offering evidence that cognitive reserve may help explain the substantial heterogeneity in memory outcomes among individuals with similar injury burdens.

### Cognitive reserve is most strongly associated with memory at lower tNAA/tCr

To our knowledge, no prior study has leveraged MRS to understand BI following IPV; the one earlier proton MRS study in this population examined post-traumatic stress [38]. Preliminary work from our group has measured metabolites by MRS in the PCG in 29 women, reporting higher glutathione and lower NAA among those with partner-inflicted BI compared to those without [39]. Our investigation therefore offers initial neurochemical evidence about memory resilience in a critical population largely absent from TBI imaging research, while complementing theoretical efforts to understand cognitive reserve [40].

Our findings suggest that the association between cognitive reserve and memory after repeated BI varies with PCG tNAA/tCr. Cognitive reserve predicted memory only when PCG tNAA/tCr, commonly interpreted as an indicator of neuronal health [33], was relatively lower (1 SD below the sample mean, a ratio of 1.29). tNAA/tCr was not significantly associated with the number of injuries, suggesting that it may not simply index cumulative exposure. This interpretation builds on a well-replicated group-level effect. A meta-analysis of 62 NAA studies found NAA reduced after TBI at every severity, though not after subconcussive hits [27], with reviews of repetitive brain trauma reporting the same direction of change [41]. PCG NAA is reduced within the first 24 hours after injury [28], and lower tNAA/tCr persists longer in athletes after a second concussion [42]. The involvement of the PCG itself is consistent with prior IPV research, in which weaker functional connectivity between the right anterior insula and both the PCG and precuneus was associated with greater BI severity and poorer learning and memory [20]. Whether lower tNAA/tCr reflects reduced neuronal-metabolic capacity or an adaptive metabolic response cannot be determined here.

The metabolic demands of memory formation offer a plausible explanation for why PCG tNAA/tCr moderated the association between cognitive reserve and memory. Experimental studies show that memory formation requires increased presynaptic mitochondrial energy production [43] and that astrocytic metabolic signaling is necessary for synaptic plasticity and recognition memory [44]. Lower PCG tNAA/tCr may therefore reflect lower metabolic capacity to meet these demands, making memory more dependent on adaptive processes such as cognitive reserve. Our findings therefore suggest a partial biological account of the episodic-memory difficulties observed in the cohort.

### Mechanistic and clinical implications

Our findings offer a key methodological implication for BI MRS, underscoring that moderation analyses may reveal information about cognitive outcomes that direct associations might otherwise fail to detect. The field has overwhelmingly tested whether metabolites differ between injured and uninjured groups or correlate with symptom burden, establishing a consistent reduction in NAA after TBI but inconsistent findings for other metabolites, moderated by region, severity and time since injury [27]. By instead testing whether metabolic state moderated the relationship between cognitive reserve and memory, we identified information that was not apparent from that metabolite’s direct association with memory.

The conditional results also have implications for study design and clinical assessment. Studies of cognitive reserve after BI have generally reported associations with cognitive performance without testing moderation [45], and where moderation has been tested, in mild TBI, the reserve effect was larger in injured participants than in community controls [46]. Moderation analyses in preclinical Alzheimer’s disease have similarly examined whether cognitive reserve modifies the relationship between brain pathology and cognition [47], placing our findings within a broader effort to understand when reserve supports cognitive resilience. A larger observational study in the CENTER-TBI cohort (*n* = 3,377), however, reported that pre-injury reserve proxies predicted outcomes largely independently of injury severity [48]. The conditions under which reserve is most strongly associated with cognitive outcomes therefore remain uncertain. Studies reporting no protective effect may have sampled participants whose exposure never reached the level at which any protective benefit of cognitive reserve becomes expressed, which may offer one explanation for the inconsistency in this literature.

In addition, our findings suggest that memory performance following BI should be assessed in relation to each individual’s own expected ability. Population norms alone describe how a woman performs relative to other people, but not how far she has moved from her own expected level. To illustrate, a woman who once performed at the 99th percentile on a memory test may, after repeated brain injuries, perform at the 50th percentile. Although cognitive reserve may have kept her performance within a normal range, it would not have prevented loss, and against population norms alone her now-average performance could go unflagged even though she has lost a great deal of memory function. As is done in practice, comparing current memory performance against an estimate of premorbid functioning improves identification of women whose harm would otherwise remain “invisible” [20,49,50], and who might benefit from cognitive rehabilitation [51]. Such idiographic approaches to neuropsychological assessment are particularly important when evaluating individuals with high premorbid functioning. Importantly, performance on standardized neuropsychological measures may not correspond closely to day-to-day functioning, and individuals with a trauma history show meaningful intra-individual and within-group variability in cognitive performance [52].

### Limitations

Several limitations merit consideration. As a cross-sectional study, we cannot establish that partner-inflicted BI caused the observed memory differences, nor can we ascertain whether neurochemical state precedes the expression of cognitive reserve. Lower tNAA/tCr may represent the metabolic condition under which cognitive reserve becomes relevant, but it could alternatively reflect pre-existing tissue health or biological processes associated with better memory. Relatedly, as we did not have longitudinal data, we could not test brain maintenance, the third concept in the consensus framework, which refers to the relative absence of change in neural resources or neuropathology over time as a determinant of preserved cognition [25]. Brain reserve and brain maintenance are separated by whether individuals differ in their initial level of neurobiological capital or in the rate at which it changes [25]. Future work should therefore incorporate longitudinal imaging to establish whether the neurochemical differences reported here reflect lower initial capacity or less successful brain maintenance. The consensus framework notes that although longitudinal designs optimally address the components of cognitive reserve, cross-sectional studies can identify variables that appear critical for cognitive reserve, establish preliminary observations, and provide insight into neurobiological mechanisms [25], which is the contribution intended here.

Our sample was restricted to women with exposure to physical IPV. We did not include women without exposure to IPV, because physical IPV carries psychiatric, social and physical consequences of its own that would be confounded with BI in such a comparison. In addition, the relatively high educational attainment of the cohort may have restricted variation and reduced sensitivity to education as an alternative proxy for cognitive reserve. Education is itself a common proxy for cognitive reserve, so adjusting for it could in principle remove variance shared with the exposure of interest, though refitting the models without education left the interaction unchanged (Supplementary Table 3). BI counts were retrospectively reported by participants whose memory may itself have been affected, although the use of two structured interviews reduces concern that one interview definition drove the findings. Injuries were sustained a median of 9.8 years before assessment among those reporting any partner-inflicted BI, so the findings describe the chronic rather than the acute phase. As a ratio, lower tNAA/tCr can reflect less NAA, more creatine, or both. After mild TBI in rats, for example, creatine fell more than NAA (44.5 vs. 29.5%), so NAA/Cr increased despite lower NAA [53]. The tNAA/tCr differences here therefore cannot be attributed to NAA alone [54]. The direction of creatine change after injury is itself uncertain. Elevated creatine can reflect increased high-energy phosphate turnover in glia responding to injury, neuroinflammation or infection [55], whereas reduced creatine has been reported in former professional football players with chronic repetitive head impact exposure [56]. Here, neither tNAA nor tCr alone moderated the association between cognitive reserve and memory. tNAA/tCr may therefore reflect neuronal integrity and energetic state jointly rather than NAA alone, and the component analyses cannot isolate either.

The study combined MRS with examiner-administered and computerized memory testing alongside structured brain-injury assessment in a population that is underrepresented in neurotrauma research and difficult to recruit. In addition, the participants who completed neuroimaging were a subset of the full cohort, which constrained statistical power and raised the possibility that the participants who were scanned differ systematically from those who were not, a known concern in neuroimaging studies of IPV [57]. Participants who underwent MRS did not differ from those who did not on age, education, premorbid functioning, injury burden, memory or post-traumatic stress symptoms, although these comparisons cannot exclude selection bias or bias from missing data.

Although the findings were largely consistent across BI thresholds, covariate specifications, memory measures, injury mechanisms (blunt force vs. strangulation), and leave-one-out analyses, replication in larger cohorts enriched for a high burden of neurotrauma is needed to establish their magnitude. The internal comparison among women who had all experienced physical IPV is also an important design strength. Participants with and without partner-inflicted BI shared exposure to violence and many accompanying psychiatric and social burdens, reducing confounding by the broader effects of IPV. At the same time, the cohort included only women who survived their injuries and were safe enough to participate. Memory impairment may itself impede study enrollment and completion, so survivors with the greatest cognitive difficulties may have been underrepresented.

## Conclusion

BI was associated with poorer memory, yet BI burden explained only part of the substantial variation among participants who had sustained ten or more BIs. Our findings offer the first behavioral and neurochemical evidence that cognitive reserve is associated with memory after partner-inflicted BI conditional on injury burden and neurochemical state. Ultimately, by showing when cognitive reserve matters, this study offers empirical evidence for an emerging account of the cognitive reserve hypothesis, and points to a biologically grounded mechanism for why, despite repeated neurotrauma, memory can remain resilient.

## Acknowledgments

We sincerely thank the women who took part in this study, and the study staff who contributed to data collection and curation. CWM was supported by graduate funds from the Knight-Hennessy Scholarship during research.

## Funding

This work was funded by grants from the National Institutes of Health (R01NS112694) (EMV). In addition, this work was conducted with support from UM1TR004408 award through Harvard Catalyst | The Harvard Clinical and Translational Science Center (National Center for Advancing Translational Sciences, National Institutes of Health) and financial contributions from Harvard University and its affiliated academic healthcare centers. The content is solely the responsibility of the authors and does not necessarily represent the official views of Harvard Catalyst, Harvard University and its affiliated academic healthcare centers, or the National Institutes of Health.

## Competing interests

CWM has no competing interests to declare. APL is employed by Mass General Brigham, receives funding from the National Institutes of Health (R01NS112694, R01NS121225), is a co-founder of Brainspec and is a consultant to Moncton MRI. SM has no competing interests to declare. AG has no competing interests to declare. EMR is on the advisory board of Brainspec (unpaid) and a consultant for Aletheia Biotherapeutics. ALJD has no competing interests to declare. SN has no competing interests to declare. EMV is employed by Massachusetts General Hospital and receives funding from the National Institutes of Health (R01NS112694).

## Data, code, and materials availability

Direct identifiers have been removed from the analysis dataset. The de-identified data supporting this study are available from the corresponding author upon reasonable request. All data will be uploaded to the Federal Interagency Traumatic Brain Injury Research (FITBIR) data repository after completion of the study, at https://fitbir.nih.gov/.

## Methods

### Experimental design

This was a cross-sectional study of community-dwelling women with lifetime exposure to at least one instance of physical intimate partner violence. The present study was designed to test whether cognitive reserve protects memory in women who have experienced partner-inflicted brain injury and under what neurobiological conditions.

### Participants

Data collection took place between 2020 and 2026 as part of a larger ongoing study of partner-violence-related brain injury [18,20]. Participants were recruited through flyers distributed by community partners, an institutional online research platform, and social media. Community partners included hospital-based programs, police-department-affiliated initiatives, and community organizations focused on violence prevention and survivor support. To be eligible, women had to have experienced at least one instance of physical partner violence and be at least 18 years old. Women were excluded from the neuroimaging arm of the study for any of the following: a brain injury within the preceding three months; a lifetime history of moderate or severe TBI; a psychiatric disorder judged likely to interfere with participation; a moderate or severe substance use disorder within the preceding six months, or a chronic history of substance use judged likely to confound results; inadequate command of English; pregnancy; a neurological disorder such as epilepsy judged likely to confound results; or any metal on or in the body posing a magnetic resonance safety risk or causing image distortion. A personal history of BI was not required. Sixty-two women form the analytic cohort. A neuroimaging subset completed spectroscopy (n = 51 to 53 depending on voxel). All processes and procedures were approved by our hospital’s Institutional Review Board and all participants provided informed consent.

### Brain injury

Exposure to BI was established through two structured interviews administered by a trained examiner. The Brain Injury Severity Assessment (BISA; [58]) applies the American Congress of Rehabilitation Medicine criteria [59] to acts specifically inflicted by an intimate partner as well as any other non-abuse related BIs in childhood and adulthood. Any event where traumatically induced disruption of brain function was evident from loss of consciousness, amnesia for the period surrounding the event, altered mental state at the time, or focal neurological signs was counted as a BI. The Ohio State University TBI Identification Method [60] assesses for injuries sustained across the lifespan from any cause, probing for loss of consciousness, feeling dazed, or a gap in memory. The number of BIs per participant was determined via findings from both interviews, and all possible brain injuries were reviewed by at least one BI expert. Consistent with these criteria and with earlier reports from this cohort, altered consciousness arising from violent shaking, and from the hypoxia and/or ischemia that strangulation produces, was counted as a BI. Where a participant described altered consciousness following a partner-inflicted event, further questioning established how the injury occurred. Events the participant attributed to strangulation or choking alone were recorded as hypoxic/ischemic; all remaining events, including those combining blunt force with strangulation and those arising from violent shaking, were grouped as blunt force for the mechanism comparison. Strangulation that did not alter consciousness (i.e., no loss of consciousness, alteration in consciousness, or post-traumatic amnesia) was not counted as a BI. BIs that resulted in a loss of consciousness for less than 30 minutes were labeled as mild, and all others as moderate-severe [59]. No partner-inflicted injury in this cohort met moderate-severe criteria (by exclusion), so injury severity could not be examined as a covariate.

Importantly, we use the term “BI” rather than TBI because partner-inflicted injuries include hypoxic-ischemic injury from strangulation, which falls outside standard definitions of TBI. TBI is retained when describing studies limited to traumatic forces to the brain.

The exposure of interest was the cumulative count of partner-inflicted BIs. Counts were severely right-skewed (median 1, interquartile range 0 to 5, range 0 to 420; adjusted Fisher-Pearson *G*1 = 7.62) with a maximum reported value of 420, and therefore log-transformed. Following prior work in this cohort [14,61], counts above 25 were Winsorized to 25, since counts above that level cannot be estimated with confidence from retrospective report. Winsorizing BI counts at 25 and log transformation reduced skew to 0.64. Among plausible exposure codings, the logarithmic specification gave the best fit by Akaike information criterion (Fig. S1). The highest-burden band of ten or more injuries was defined following prior work in this cohort [14].

### Cognitive assessment

Memory and learning were assessed by a trained psychometrician using the California Verbal Learning Test, Third Edition (CVLT-3) [62]. The CVLT-3 was administered remotely by video call (Zoom). Participants were read a list of 16 words five times and asked to recall as many words as possible immediately following each reading, and again after a 20-minute delay. Words recalled across trials 1 to 5, expressed as a scaled score, served as a measure of learning, whereas the words recalled after the delay, long-delay free recall (LDFR), served as a measure of memory. LDFR is reported as a scaled score with a normative mean of 10 and standard deviation of 3, and served as the primary outcome. An episodic composite averaging the learning and long-delay recall scores, which share a common scaled-score metric, was examined as a secondary outcome to establish that findings did not rest on a single metric.

Participants also independently completed CNS Vital Signs, a computerized battery with age-adjusted norms [63]. Fifteen domains were available in this cohort and were classified as memory (composite, verbal, visual and working memory) or non-memory (complex attention, cognitive flexibility, executive function, processing speed, psychomotor speed, motor speed, reaction time, simple attention, sustained attention, reasoning and social acuity) for the specificity analysis; the subtests contributing to each domain are listed and described in Supplementary Tables 9 and 10. Every domain score returned by the battery was analyzed, with performance validity established from the dedicated effort measures described below rather than from any single instrument, except that standard scores more than four standard deviations from the normative mean (below 40 or above 160) were excluded as implausible; these included off-scale values the report prints for very poor performance, and removed 33 of 930 domain scores, none in a memory domain. CNS Vital Signs was administered remotely in a private location of the participant’s choosing.

Cognitive reserve was estimated using the Test of Premorbid Functioning (ToPF) [64], a single-word reading measure. Performance on the ToPF, like that on other single-word reading tests, is relatively resistant to change after BI. Word-reading scores are stable over the year following mild injury, with instability confined to moderate and severe injury [65], and ToPF performance does not differ across injury severity groups [49]. Lifetime moderate or severe TBI was an exclusion criterion here. The ToPF was therefore used as an estimate of premorbid functioning and a proxy for cognitive reserve. Scores are reported as standard scores with a normative mean of 100 and standard deviation of 15. We used the ToPF standard score derived from word-reading performance, not the demographically predicted score. Cognitive reserve cannot be measured directly [22], so this score is a proxy; for brevity, this is referred to as cognitive reserve in the Results.

Performance validity was assessed with the Test of Memory Malingering [66] and two embedded validity measures: reliable digit span from the Wechsler Adult Intelligence Scale, Fourth Edition digit span forward and backward trials, with a score of 7 or above treated as a pass [67,68] and CVLT-3 forced-choice recognition, with a score of 15 or above treated as a pass [62]. Performance was deemed valid based on clinical judgment and if at least two of the three measures were valid. Analyses in the main text retain all participants with valid data on the relevant measures. Pass rates for individual tests are reported in Supplementary Table 11, and a sensitivity analysis omitting participants flagged on any indicator is reported in Supplementary Table 12. Three participants were separately identified by the study assessor as warranting review; repeating every analysis without them is reported in Supplementary Table 12.

### Covariates

Years of education and post-traumatic stress symptom severity measured with the PTSD Checklist for DSM-5 (PCL-5) [69] were included as covariates in all adjusted models. Age was not adjusted for in models of the memory measures, because those scores are already age-corrected against published norms, but was retained in the spectroscopy models, where metabolite concentrations vary with age and age would otherwise confound the predictor. Post-traumatic stress was also adjusted for because PTSD is associated with poorer performance across cognitive domains, with the largest effects in verbal learning and memory [70], and with abnormalities in the neurochemical systems sampled by spectroscopy [71]. Relatedly, probable PTSD (PCL-5 ≥ 33) was present in 25 of 62 participants and was unevenly distributed across exposure bands, affecting 9 of 12 participants (75%) in the highest-burden group against 40% of those reporting no partner-inflicted injury and 27% of those reporting one to nine (*χ*^2^ = 8.32, *P* = .015). The relationship was not monotonic, however, and continuous symptom severity was associated with neither injury burden (*ρ* = 0.08, *P* = .52) nor delayed recall (*ρ* = 0.03, *P* = .81), so the adjustment guards against a plausible confounder rather than removing variance that carries the effect. Additional variables examined in sensitivity analyses were depression symptoms with the Patient Health Questionnaire-9 [72], alcohol use with the Alcohol Use Disorders Identification Test [32], childhood maltreatment with the Childhood Trauma Questionnaire [73], abuse severity with the Composite Abuse Scale [74], race and ethnicity, and brain injuries from childhood and from non-partner sources in adulthood, entered as separate covariates following the approach used previously in this cohort [75], body mass index derived from the height and weight recorded at telephone screening, and current medication use, entered in separate models as any medication (yes/no) and as psychotropic medication (yes/no).

### Magnetic resonance spectroscopy

Magnetic resonance spectroscopy (MRS) was acquired on a Siemens MAGNETOM Prisma 3T scanner with a 32-channel head coil at the Athinoula A. Martinos Center for Biomedical Imaging. Localizer images were acquired using T1-weighted Multi-echo MPRAGE (TE1 1.81 ms, TE2 3.6 ms, TE3 5.39 ms, TE4 7.18 ms, TR 2500.0 ms, 0.8×0.8×0.8 mm isotropic resolution, PAT mode GRAPPA: Accel. Factor 2). Voxels were placed in the posterior cingulate gyrus, dorsolateral prefrontal cortex, and corona radiata (Fig. 4). In each voxel, MRS was acquired using point-resolved spectroscopy with repetition/echo time of 1700/30 ms, 128 averages, 2×2×2 cm^3^, bandwidth of 4000 Hz, and 2078 data points. An automated shim was used, and manual shimming was used if linewidths exceeded 15 Hz. An unsuppressed water reference was also acquired. MRS data were pre-processed using Osprey (v.2.5.0) [76] and post-processed using LCModel [77] using the default basis set included with Osprey. Metabolites were quantified using the quantification and tissue correction built into Osprey; as all analyses used ratios to total creatine, tissue-composition corrections cancel in the reported measures. Quality assurance was conducted using automated and manual assessment [78]. Spectra with major artifacts or poor data quality were excluded (two from the DLPFC voxel and four from the corona radiata), leaving 51, 50 and 49 spectra in the posterior cingulate, dorsolateral prefrontal and corona radiata voxels; the same four participants lacked delayed-recall scores, so 47, 46 and 45 contributed to the adjusted models. All MRS methods are summarized in Supplementary Table 13 [79,80].

**Fig. 4.**
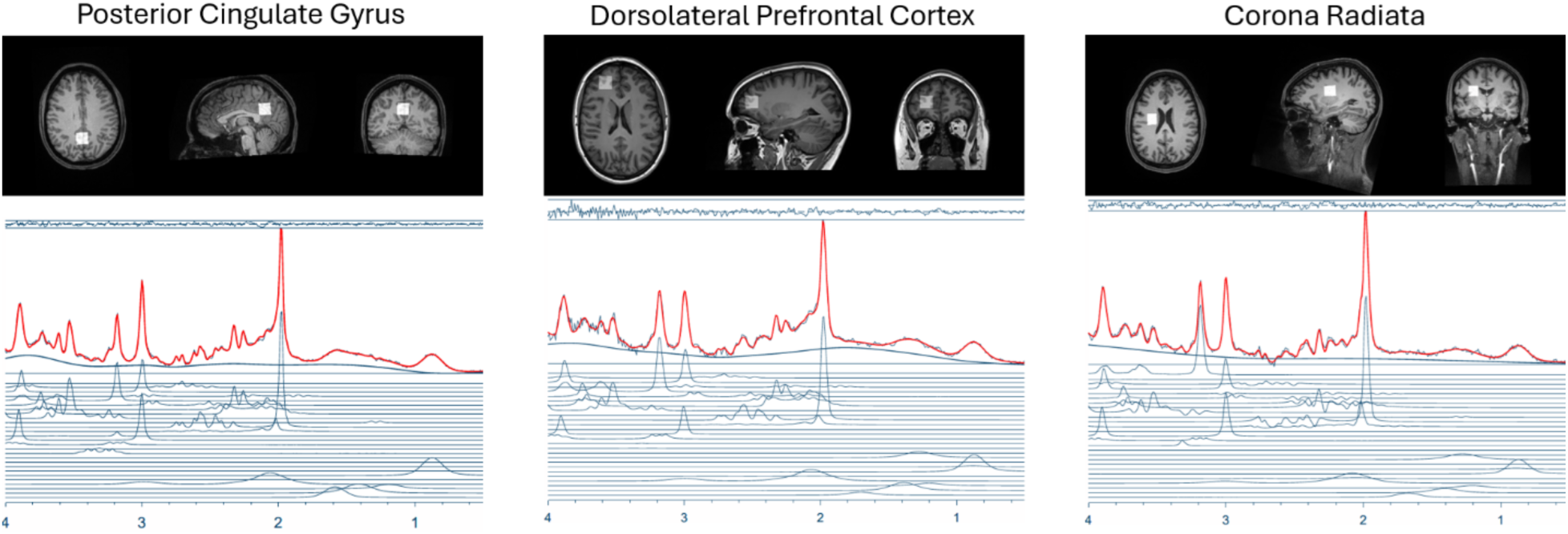
Representative voxel location and spectra for the posterior cingulate gyrus (left), dorsolateral prefrontal cortex (middle), and corona radiata (right).

### Statistical analysis

All continuous variables were standardized before modeling, so regression coefficients are in standard deviation units and interaction coefficients represent differences in slope. Models were estimated by ordinary least squares with heteroskedasticity-consistent HC3 standard errors [81], which perform well at the sample sizes used here [82]. Inference used the t distribution with residual degrees of freedom. Estimates are interpreted by magnitude and interval width rather than by whether they cross a significance threshold, since a difference between a significant and a non-significant result is not itself evidence of a difference [83]. Sample sizes differ across analyses because not every participant completed every measure. Each model accordingly uses all participants with complete data on its own variables. Associations within a single exposure band are reported as unadjusted Spearman correlations, because the highest-burden band contains twelve participants, with analytic n varying by measure, and cannot support the covariate-adjusted models used elsewhere. Inferential claims rest on the interaction models, which are fit in the full cohort and adjusted.

Subgroup correlations used Spearman rank coefficients, which are insensitive to the extreme injury counts present in the cohort. 95% confidence intervals for subgroup correlations were obtained by Fisher z-transformation (Fig. 2B); bias-corrected and accelerated bootstrap with 10,000 resamples was used for the robustness and mediation analyses. Conditional effects of cognitive reserve were estimated at one standard deviation above and below the moderator mean [84]. The influence of individual participants was assessed by refitting each model with each participant omitted in turn, and out-of-sample predictive value was assessed by leave-one-out cross-validation, comparing the cross-validated R^2^ of models with and without the interaction term.

Multiple comparisons were controlled by the Benjamini-Hochberg false discovery rate [85] within analysis families. Total NAA to creatine was the metabolite of interest, both because it is the established marker of neuronal integrity and because its resonance is the largest in the proton spectrum and therefore the most reliably quantified, so correction was applied across the three voxels in which it was measured. The nine remaining creatine-referenced metabolite and voxel combinations were treated as a separate family. The reserve-by-injury and reserve-by-tNAA/tCr interactions were the primary tests; analyses of the remaining metabolites, alternative thresholds, cognitive domains and injury mechanisms are exploratory.

Finally, we tested whether the metabolite might mediate, rather than moderate, the association between injury and memory. Injury burden did not predict posterior cingulate tNAA/tCr (a path, *β* = 0.02, *P* = .90), and the metabolite did not predict memory once injury was in the model (b path, *β* = −0.20, *P* = .22). The indirect effect was compatible with zero (*a* × *b* = −0.003, 95% bootstrap CI −0.09 to 0.04), while the direct effect of injury on memory was unchanged (*β* = −0.50, *P* = .002). Moderation and mediation are not mutually exclusive, and this null result does not rule out an indirect pathway or establish that the metabolite is unrelated to accumulated injury.

### Effect sizes

Effect sizes for correlational analyses are reported as Spearman rank correlation coefficients, chosen because injury count is severely right-skewed. For all regression models, effect sizes are reported as standardized regression coefficients, defined as the change in standard deviations of the outcome for a one standard deviation change in the predictor while holding all other predictors constant. All continuous variables were standardized before modeling, so coefficients are directly comparable across models and across outcomes measured on different scales.

Moderation was specified as a product term between cognitive reserve and the candidate moderator, with both constituent terms retained in the model:

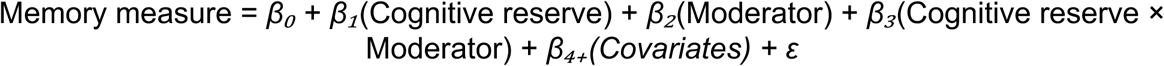

The coefficient of interest is *β*₃, which gives the difference in the reserve-to-memory slope per unit change in the moderator. Simple slopes at specific moderator values were computed as *β*₁ + *β*₃(value), with standard errors derived from the full covariance matrix rather than by combining the component standard errors, and were tested against the residual degrees of freedom of the fit model. Analyses were conducted in Python 3.12 and GraphPad Prism 11.0.2.

## Supplementary Information

**Fig. S1.**
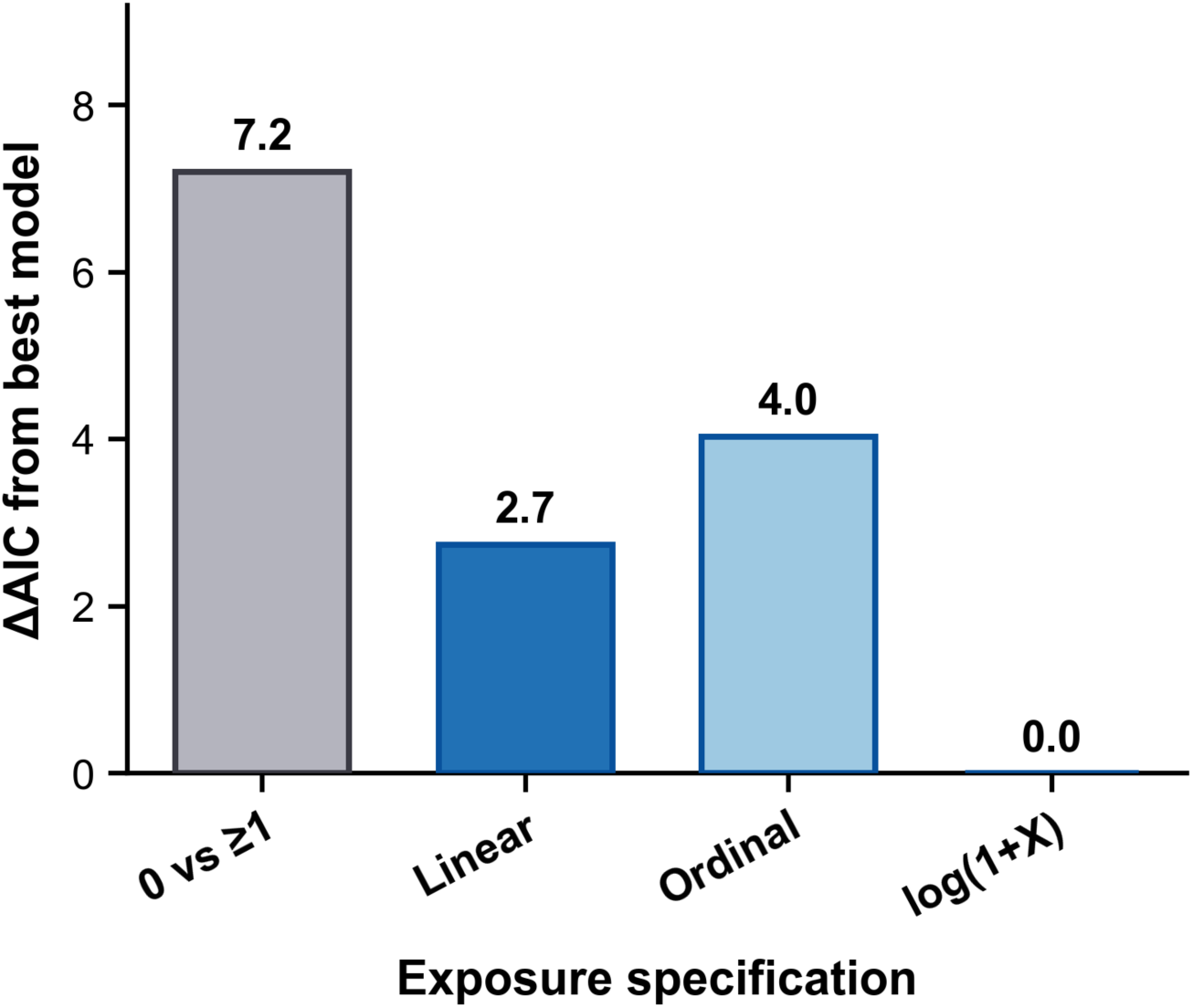
Model fit under alternative codings of injury exposure. Bars give the difference in Akaike information criterion between each coding of partner-inflicted injury and the best-fitting coding, in models of delayed verbal recall adjusted for education and PTSD, with counts capped at 25 as in the primary analysis. Lower values indicate better fit, and a difference below about two units is conventionally taken as no meaningful difference between models. The logarithmic coding, used for the association between injury burden and memory and as a covariate in the spectroscopy models, fit best, whereas the ordinal, linear-count and binary any-injury codings each fit worse by more than two units. Sensitivity analyses provided in Supplementary Table 3 show that the associations reported here do not depend on model choice. Notably, the reserve-by-injury interaction was not significant under any continuous coding of injury burden, including the raw count, its logarithm, its square root and its rank (all *P* ≥ .40), indicating that the interaction reflects a contrast between exposure bands rather than a graded function of injury number.

**Fig. S2.**
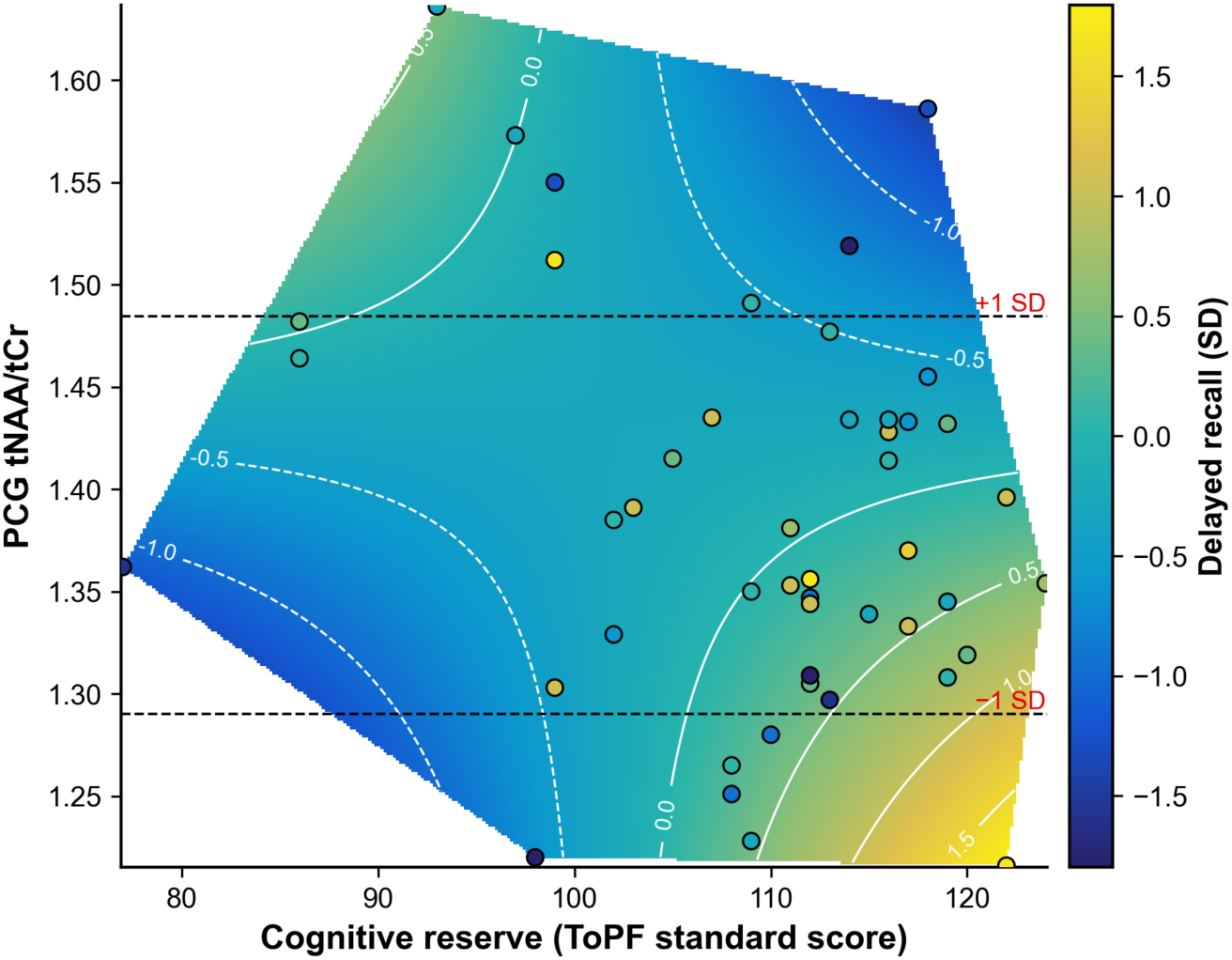
Predicted delayed recall across the observed joint distribution of cognitive reserve and PCG tNAA/tCr. Shading and white contours show delayed recall (SD units) predicted by the reserve-by-tNAA/tCr model with covariates held at their means, drawn only within the convex hull of the observed data so that no values are extrapolated. Points are participants (*n* = 47), colored by observed delayed recall. Dashed lines mark tNAA/tCr 1 SD below (1.29) and above (1.48) the mean. The association between cognitive reserve and recall is strongest at lower tNAA/tCr, and predicted values cross within the observed range, indicating a crossover interaction. Estimates in the sparsely sampled corners rest on few participants.

---

**Supplementary Table 1.**
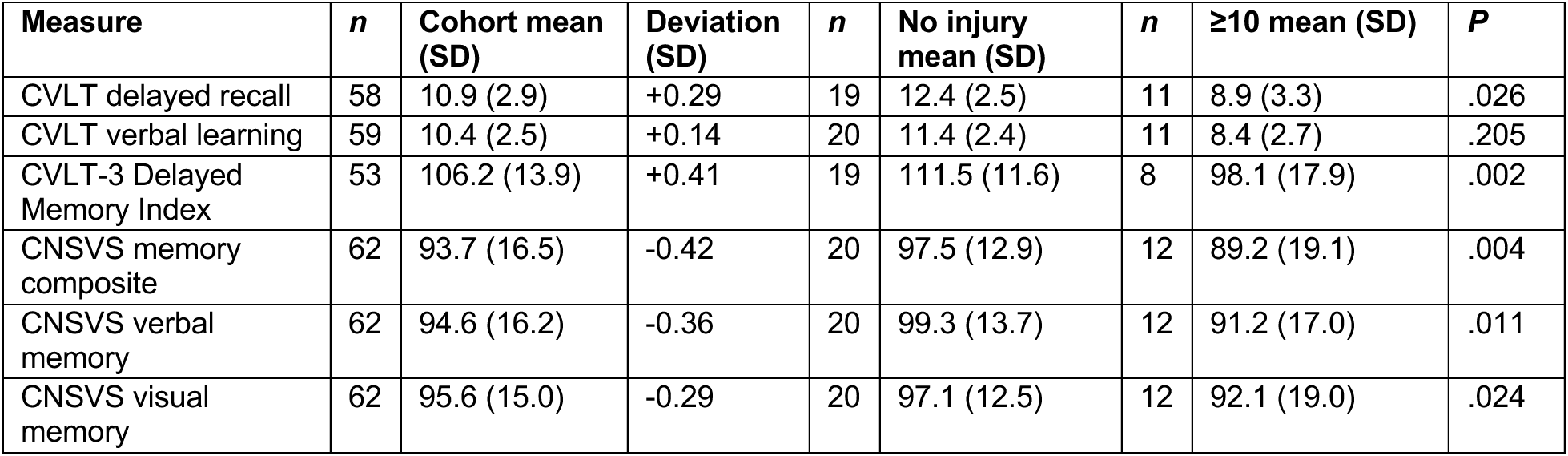
Figure 1 **memory performance.** Scores are given for the whole cohort and for the two exposure groups displayed in Figure 1, so that the distributions shown graphically can be read as numbers. Deviation is expressed in normative standard deviation units, where zero is the performance expected of the general population. *P* values are one-sample *t* tests of the whole-cohort mean against the published normative mean and are uncorrected. Cohort education and premorbid functioning exceed those of the normative samples, so deviations from published norms should not be read as injury effects.

| **Measure** | ***n*** | **Cohort mean (SD)** | **Deviation (SD)** | ***n*** | **No injury mean (SD)** | ***n*** | **≥10 mean (SD)** | ***P*** |
|---|---|---|---|---|---|---|---|---|
| CVLT delayed recall | 58 | 10.9 (2.9) | +0.29 | 19 | 12.4 (2.5) | 11 | 8.9 (3.3) | .026 |
| CVLT verbal learning | 59 | 10.4 (2.5) | +0.14 | 20 | 11.4 (2.4) | 11 | 8.4 (2.7) | .205 |
| CVLT-3 Delayed Memory Index | 53 | 106.2 (13.9) | +0.41 | 19 | 111.5 (11.6) | 8 | 98.1 (17.9) | .002 |
| CNSVS memory composite | 62 | 93.7 (16.5) | -0.42 | 20 | 97.5 (12.9) | 12 | 89.2 (19.1) | .004 |
| CNSVS verbal memory | 62 | 94.6 (16.2) | -0.36 | 20 | 99.3 (13.7) | 12 | 91.2 (17.0) | .011 |
| CNSVS visual memory | 62 | 95.6 (15.0) | -0.29 | 20 | 97.1 (12.5) | 12 | 92.1 (19.0) | .024 |

**Supplementary Table 2.**
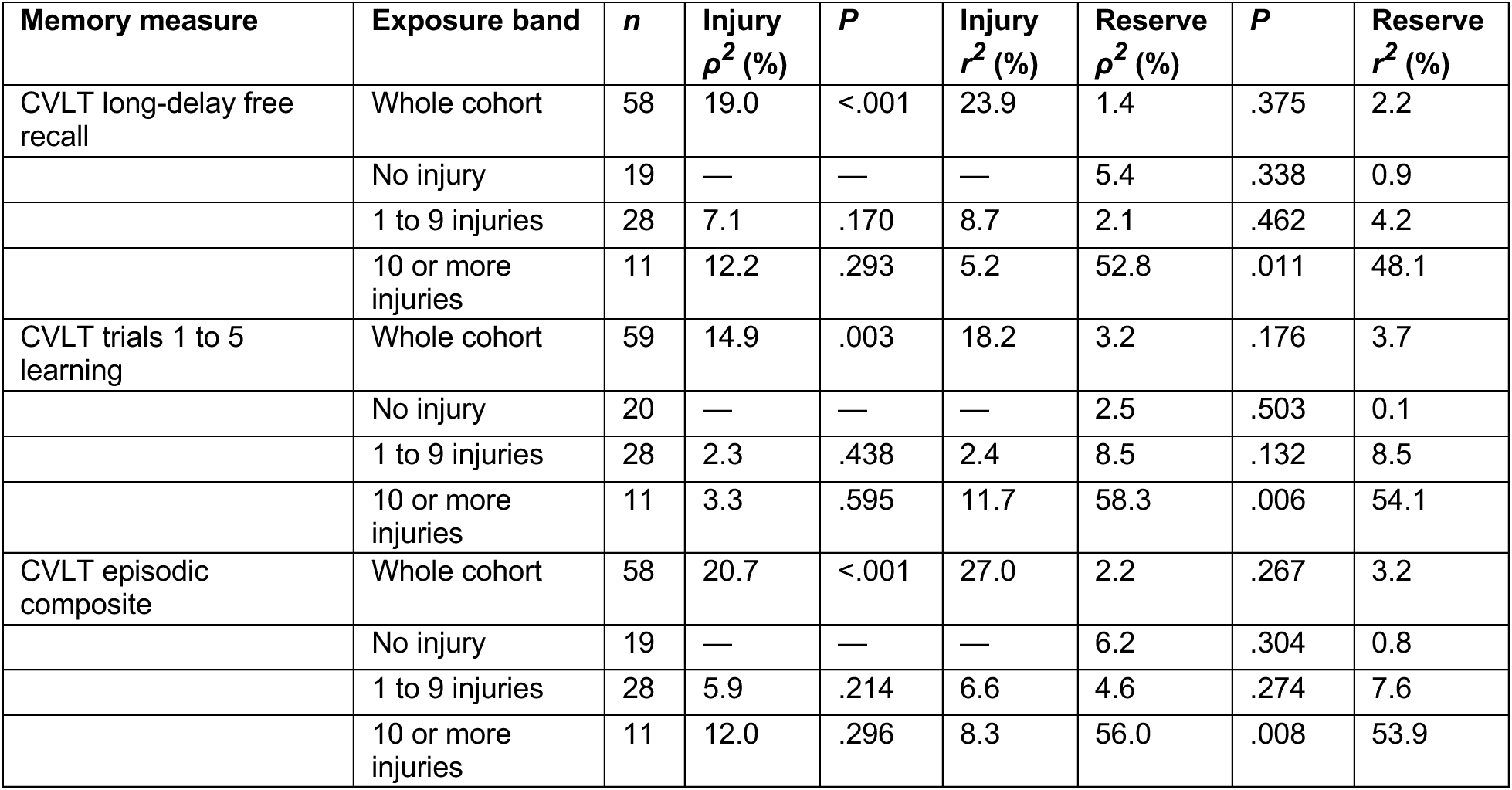
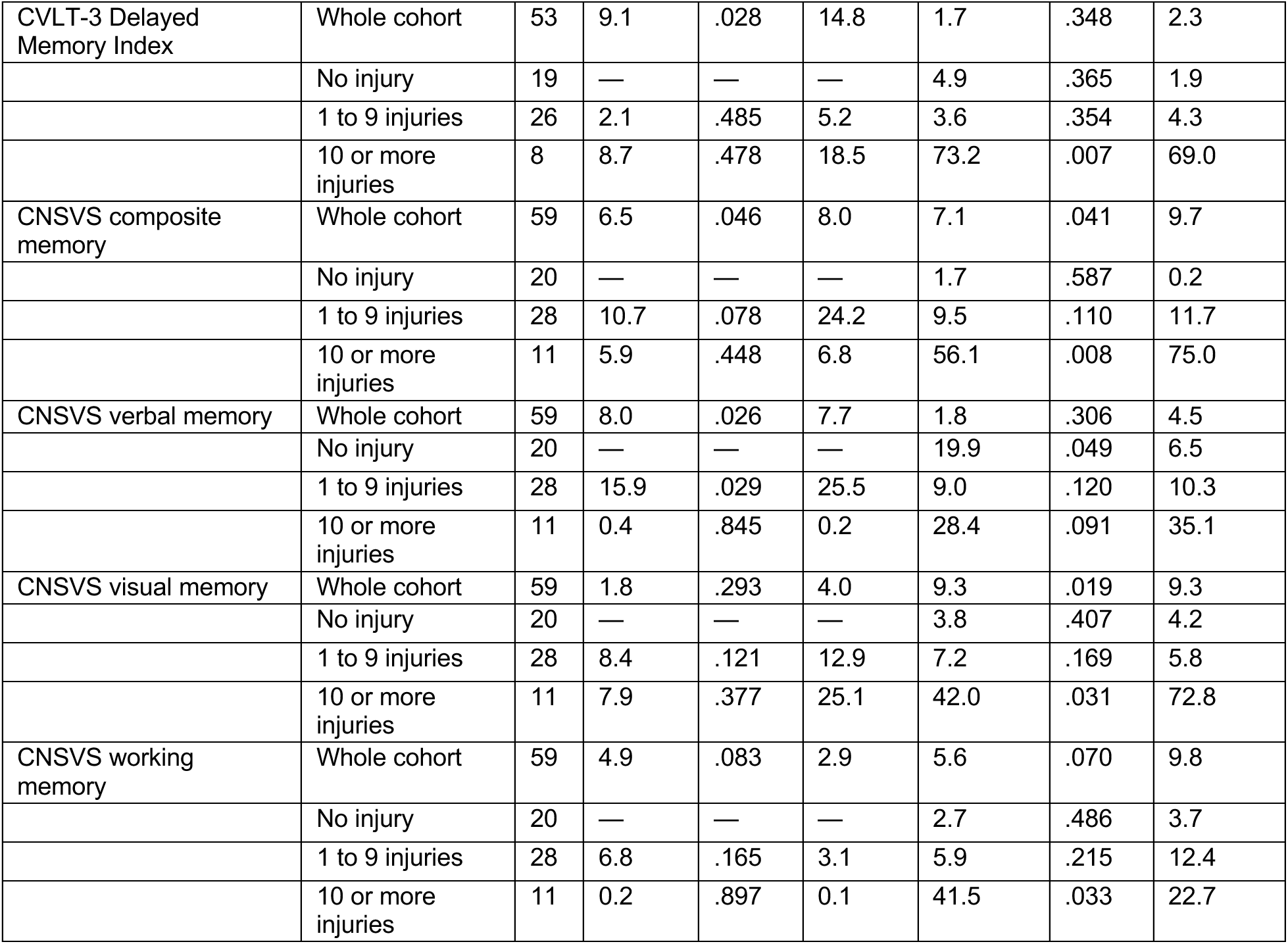
Variance in memory explained by injury burden and by cognitive reserve, within each exposure band. For each memory measure, *ρ^2^* is the squared Spearman correlation, expressed as a percentage, with either cumulative partner-inflicted brain injury or premorbid functioning, and *r^2^* is the corresponding squared Pearson correlation, the proportion of variance explained in the untransformed scores. Given that several within-band subsamples are small and are therefore sensitive to individual observations, *P* values are reported for the Spearman correlation. To determine where each predictor carries explanatory weight, associations are given for the whole cohort and separately within each exposure band. For instance, among participants reporting ten or more injuries, premorbid functioning is strongly associated with long-delay free recall (*ρ*^2^ = 52.8%, *r*^2^= 48.1%) whereas injury burden is only weakly associated (*ρ*^2^ = 12.2%, *r*^2^ = 5.2%). Injury burden is entered as log(1 + count) for the whole cohort and as the raw count within bands, and is undefined for participants reporting no injury. These are unadjusted associations, reported to show the relative contribution of each predictor rather than as the inferential models. Bands with fewer than ten participants on a given measure are shown for completeness but sample size should be considered.

| **Memory measure** | **Exposure band** | ***n*** | **Injury *ρ*<sup>*2*</sup> (%)** | ***P*** | **Injury *r*<sup>*2*</sup> (%)** | **Reserve *ρ*<sup>*2*</sup> (%)** | ***P*** | **Reserve *r*<sup>*2*</sup> (%)** |
|---|---|---|---|---|---|---|---|---|
| CVLT long-delay free recall | Whole cohort | 58 | 19.0 | <.001 | 23.9 | 1.4 | .375 | 2.2 |
|  | No injury | 19 | — | — | — | 5.4 | .338 | 0.9 |
|  | 1 to 9 injuries | 28 | 7.1 | .170 | 8.7 | 2.1 | .462 | 4.2 |
|  | 10 or more injuries | 11 | 12.2 | .293 | 5.2 | 52.8 | .011 | 48.1 |
| CVLT trials 1 to 5 learning | Whole cohort | 59 | 14.9 | .003 | 18.2 | 3.2 | .176 | 3.7 |
|  | No injury | 20 | — | — | — | 2.5 | .503 | 0.1 |
|  | 1 to 9 injuries | 28 | 2.3 | .438 | 2.4 | 8.5 | .132 | 8.5 |
|  | 10 or more injuries | 11 | 3.3 | .595 | 11.7 | 58.3 | .006 | 54.1 |
| CVLT episodic composite | Whole cohort | 58 | 20.7 | <.001 | 27.0 | 2.2 | .267 | 3.2 |
|  | No injury | 19 | — | — | — | 6.2 | .304 | 0.8 |
|  | 1 to 9 injuries | 28 | 5.9 | .214 | 6.6 | 4.6 | .274 | 7.6 |
|  | 10 or more injuries | 11 | 12.0 | .296 | 8.3 | 56.0 | .008 | 53.9 |
| CVLT-3 Delayed Memory Index | Whole cohort | 53 | 9.1 | .028 | 14.8 | 1.7 | .348 | 2.3 |
|  | No injury | 19 | — | — | — | 4.9 | .365 | 1.9 |
|  | 1 to 9 injuries | 26 | 2.1 | .485 | 5.2 | 3.6 | .354 | 4.3 |
|  | 10 or more injuries | 8 | 8.7 | .478 | 18.5 | 73.2 | .007 | 69.0 |
| CNSVS composite memory | Whole cohort | 59 | 6.5 | .046 | 8.0 | 7.1 | .041 | 9.7 |
|  | No injury | 20 | — | — | — | 1.7 | .587 | 0.2 |
|  | 1 to 9 injuries | 28 | 10.7 | .078 | 24.2 | 9.5 | .110 | 11.7 |
|  | 10 or more injuries | 11 | 5.9 | .448 | 6.8 | 56.1 | .008 | 75.0 |
| CNSVS verbal memory | Whole cohort | 59 | 8.0 | .026 | 7.7 | 1.8 | .306 | 4.5 |
|  | No injury | 20 | — | — | — | 19.9 | .049 | 6.5 |
|  | 1 to 9 injuries | 28 | 15.9 | .029 | 25.5 | 9.0 | .120 | 10.3 |
|  | 10 or more injuries | 11 | 0.4 | .845 | 0.2 | 28.4 | .091 | 35.1 |
| CNSVS visual memory | Whole cohort | 59 | 1.8 | .293 | 4.0 | 9.3 | .019 | 9.3 |
|  | No injury | 20 | — | — | — | 3.8 | .407 | 4.2 |
|  | 1 to 9 injuries | 28 | 8.4 | .121 | 12.9 | 7.2 | .169 | 5.8 |
|  | 10 or more injuries | 11 | 7.9 | .377 | 25.1 | 42.0 | .031 | 72.8 |
| CNSVS working memory | Whole cohort | 59 | 4.9 | .083 | 2.9 | 5.6 | .070 | 9.8 |
|  | No injury | 20 | — | — | — | 2.7 | .486 | 3.7 |
|  | 1 to 9 injuries | 28 | 6.8 | .165 | 3.1 | 5.9 | .215 | 12.4 |
|  | 10 or more injuries | 11 | 0.2 | .897 | 0.1 | 41.5 | .033 | 22.7 |

**Supplementary Table 3.**
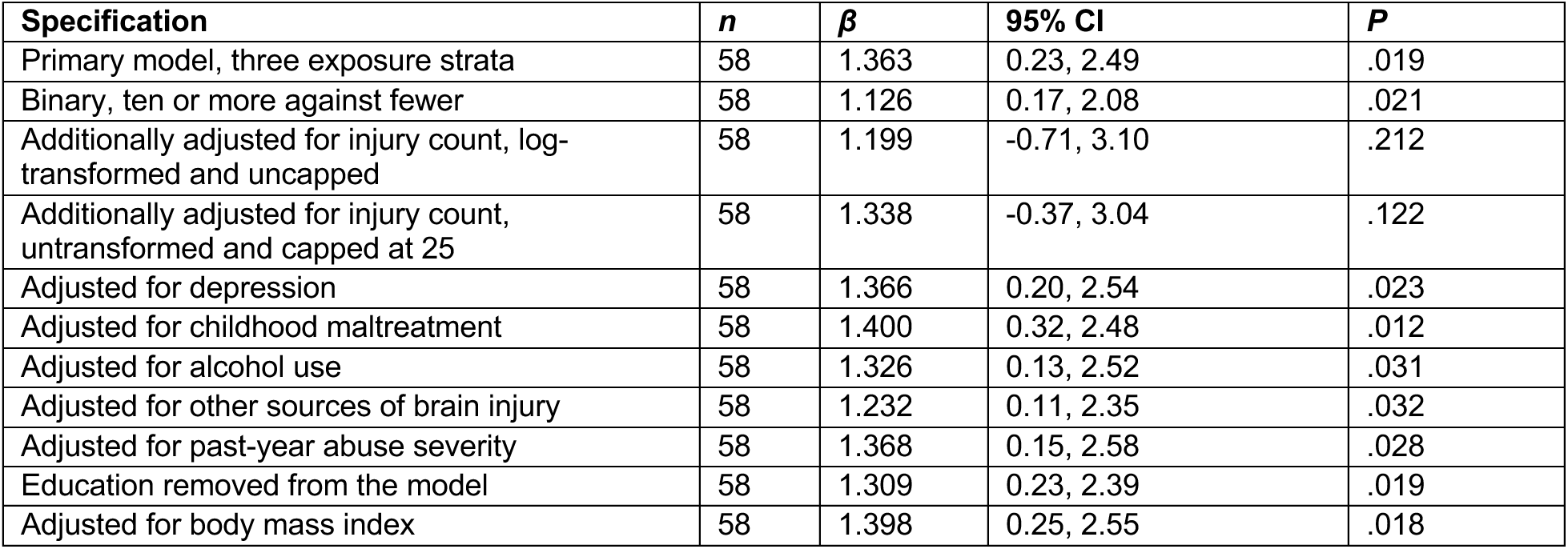
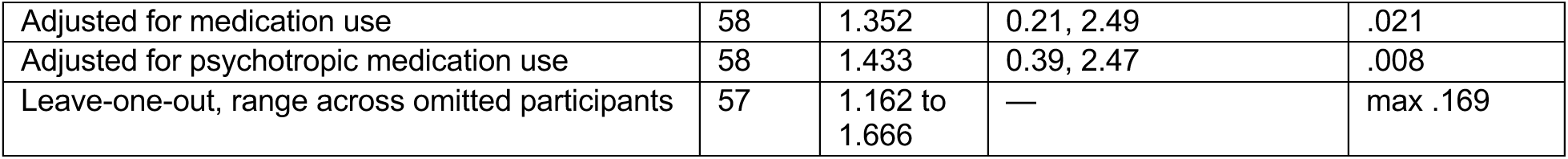
Sensitivity analyses for the reserve-by-injury interaction on delayed recall. To determine if the primary interaction depends on modeling choices, individual participants, and on the handling of extreme injury counts, we re-estimated our primary interaction under each alternative specification. The interaction rests on a small highest-burden group in which a few participants carry high leverage; we accordingly tested the interaction using the Freedman–Lane permutation method. The interaction remained significant (*P* = .024). Leave-one-out estimates refit the primary model with each participant omitted in turn; the estimate remained between 1.16 and 1.67 but was no longer significant when any one of four participants was omitted (maximum *P* = .17). Education is itself a common proxy for cognitive reserve, so adjusting for it risks removing variance the exposure of interest shares with the outcome. The final row removes education, and the interaction remains unchanged.

**Supplementary Table 4.**
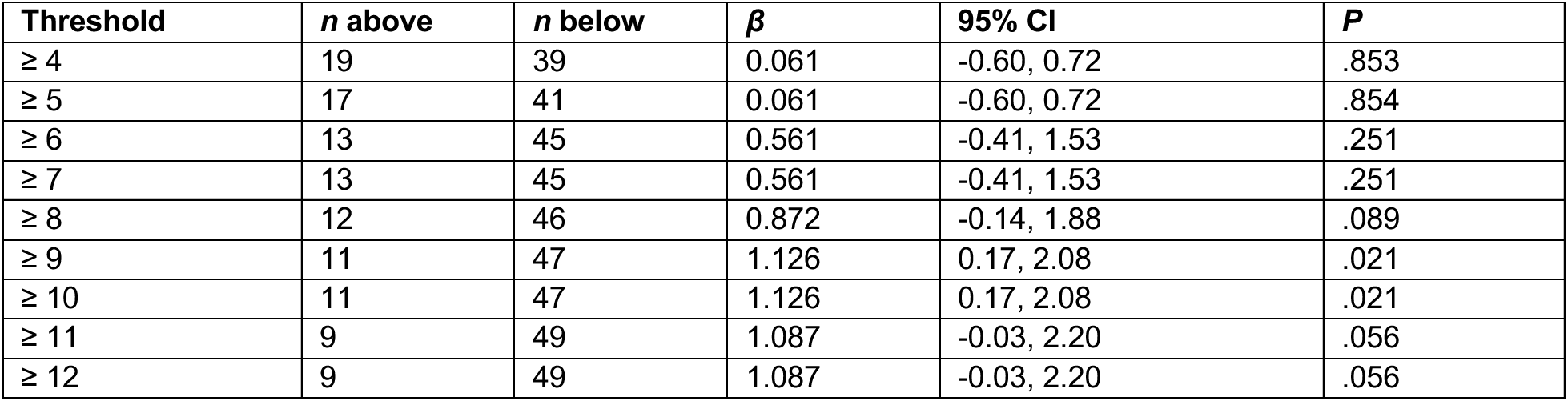
Reserve-by-injury interaction on delayed recall at every exposure threshold. To establish the exposure level at which the interaction becomes detectable, and to determine whether the finding depends on the cut-off chosen for the main analysis, we re-estimated the interaction at every candidate threshold. No participant reported exactly 6, 9 or 11 injuries, so adjacent thresholds define identical groups. All models adjust for education and post-traumatic stress symptoms with HC3 robust standard errors.

| **Threshold** | ***n* above** | ***n* below** | ***β*** | **95% CI** | ***P*** |
|---|---|---|---|---|---|
| ≥ 4 | 19 | 39 | 0.061 | -0.60, 0.72 | .853 |
| ≥ 5 | 17 | 41 | 0.061 | -0.60, 0.72 | .854 |
| ≥ 6 | 13 | 45 | 0.561 | -0.41, 1.53 | .251 |
| ≥ 7 | 13 | 45 | 0.561 | -0.41, 1.53 | .251 |
| ≥ 8 | 12 | 46 | 0.872 | -0.14, 1.88 | .089 |
| ≥ 9 | 11 | 47 | 1.126 | 0.17, 2.08 | .021 |
| ≥ 10 | 11 | 47 | 1.126 | 0.17, 2.08 | .021 |
| ≥ 11 | 9 | 49 | 1.087 | -0.03, 2.20 | .056 |
| ≥ 12 | 9 | 49 | 1.087 | -0.03, 2.20 | .056 |

**Supplementary Table 5.** The reserve-by-injury interaction does not differ detectably by injury mechanism. Partner-inflicted brain injuries were grouped by mechanism. The strangulation category contains events attributed to strangulation alone (68 events). The blunt force category contains events attributed to force applied to the head (608 events) together with events combining blunt force and strangulation (14 events) and events arising from violent shaking (14 events), because these could not be assigned to a single route of injury. Mechanism is tested on the continuous counts rather than by dichotomizing them. Both mechanism counts, log-transformed and capped as in the main analysis, were entered in a single model together with their interactions with cognitive reserve, and the two interaction coefficients were compared by a formal contrast. The two mechanisms are not collinear among injured participants (Spearman *ρ* = −0.46), so the contrast is estimable. A further row follows the approach taken in earlier work on this cohort, entering strangulation as ever against never while adjusting for total injury burden. No specification detects a difference between mechanisms. The sample cannot resolve a moderate difference, however. Thirteen participants reported any strangulation, the median count among them was two, and only one participant reported ten or more, so the exposure range over which the reserve-by-injury interaction is expressed is populated almost entirely by blunt-force injury. All models adjust for education and post-traumatic stress symptoms.

| **Specification** | ***n*** | ***β*** | **95% CI** | ***P*** |
|---|---|---|---|---|
| Blunt force, both mechanisms modeled | 58 | 0.261 | -0.13, 0.65 | .188 |
| Strangulation, both mechanisms modeled | 58 | 0.000 | -0.53, 0.53 | 1.000 |
| Difference between mechanisms | 58 | 0.262 | -0.27, 0.80 | .330 |
| Strangulation ever, adjusting for total burden | 58 | -0.035 | -0.71, 0.64 | .918 |

**Supplementary Table 6.** Reserve-by-injury interaction (ten or more BIs against fewer than ten) across all CNS Vital Signs domains. For completeness, the interaction is reported for every domain of the battery; the main text focuses on the memory domains and two comparison domains (processing speed and executive function). *q* values are Benjamini-Hochberg false discovery rates computed within the memory and non-memory families separately. A significant interaction in memory domains alongside no significant interaction elsewhere does not by itself establish that the two differ. To test the difference directly, we entered all fifteen domains into a single linear mixed model with domain family as a factor and a random intercept for participant, standardizing each domain score within itself, across all available domain observations from the 59 participants. The three-way term between cognitive reserve, injury burden and domain family was significant (*β* = 1.12, 95% CI [0.68, 1.57], *P* < .001), with the reserve-by-injury slope at 0.90 in memory domains and −0.22 in non-memory domains.

| **Family** | **Domain** | ***n*** | ***β*** | **95% CI** | ***P*** | ***q*** |
|---|---|---|---|---|---|---|
| Memory | Composite Memory | 59 | 1.298 | 0.03, 2.56 | .045 | .090 |
| Memory | Verbal Memory | 59 | 0.829 | -0.79, 2.44 | .308 | .410 |
| Memory | Visual Memory | 59 | 1.380 | 0.64, 2.12 | <.001 | .002 |
| Memory | Working Memory | 59 | 0.350 | -0.61, 1.31 | .467 | .467 |
| Non-memory | Cognitive Flexibility | 57 | -0.022 | -1.83, 1.78 | .980 | .980 |
| Non-memory | Complex Attention | 50 | -0.809 | -2.45, 0.84 | .328 | .980 |
| Non-memory | Executive Function | 58 | -0.196 | -1.90, 1.50 | .818 | .980 |
| Non-memory | Motor Speed | 57 | -0.025 | -1.28, 1.23 | .968 | .980 |
| Non-memory | Processing Speed | 58 | -0.192 | -1.00, 0.62 | .635 | .980 |
| Non-memory | Psychomotor Speed | 57 | -0.104 | -1.18, 0.97 | .847 | .980 |
| Non-memory | Reaction Time | 59 | -0.403 | -1.58, 0.77 | .495 | .980 |
| Non-memory | Reasoning | 59 | 0.155 | -0.79, 1.10 | .744 | .980 |
| Non-memory | Simple Attention | 50 | -1.790 | -5.88, 2.30 | .383 | .980 |
| Non-memory | Sustained Attention | 55 | 0.398 | -0.74, 1.53 | .484 | .980 |
| Non-memory | Social Acuity | 57 | 0.076 | -0.84, 0.99 | .869 | .980 |

**Supplementary Table 7.** Complete metabolite grid with reserve-by-metabolite interactions on delayed recall. The complete grid is given so that the primary tNAA/tCr results can be read against every comparison performed, and so that the magnitude and precision of the effects that did not survive correction remain visible. All models adjust for injury burden, age, education and post-traumatic stress symptoms. Total N-acetylaspartate to creatine was the metabolite of interest on measurement and theoretical grounds, so *q* values for the three voxels in which it was measured are Benjamini-Hochberg false discovery rates within that family, and *q* values for the nine remaining cells are computed within a separate family. Six spectra were excluded at quality control, two from the DLPFC voxel and four from the corona radiata, and four participants lacked delayed recall, so the adjusted models include 47 participants for the PCG, 46 for the DLPFC and 45 for the corona radiata. Across the analyzed samples, tNAA/tCr (mean, SD) was 1.39 (0.10) in the PCG, 1.52 (0.15) in the DLPFC and 1.61 (0.19) in the corona radiata; in the PCG, one standard deviation below and above the mean corresponds to 1.29 and 1.48. tNAA/tCr was approximately normally distributed in each voxel (Shapiro–Wilk *P* = .46, .62 and .15 for the PCG, DLPFC and corona radiata).

| **Voxel** | **Metabolite** | ***n*** | ***β*** | **95% CI** | ***P*** | ***q*, family** |
|---|---|---|---|---|---|---|
| PCG | tNAA/tCr | 47 | -0.456 | -0.71, -0.20 | .001 | .002 |
| PCG | Glx | 47 | -0.234 | -0.48, 0.01 | .058 | .284 |
| PCG | mI | 47 | -0.036 | -0.33, 0.25 | .801 | .901 |
| PCG | Cho | 47 | 0.005 | -0.43, 0.44 | .982 | .982 |
| DLPFC | tNAA/tCr | 46 | -0.347 | -0.65, -0.05 | .024 | .036 |
| DLPFC | Glx | 46 | -0.186 | -0.79, 0.42 | .539 | .901 |
| DLPFC | mI | 46 | 0.179 | -0.10, 0.46 | .203 | .507 |
| DLPFC | Cho | 46 | -0.077 | -0.58, 0.43 | .758 | .901 |
| Corona radiata | tNAA/tCr | 45 | -0.175 | -0.55, 0.20 | .345 | .345 |
| Corona radiata | Glx | 45 | 0.103 | -0.35, 0.56 | .650 | .901 |
| Corona radiata | mI | 45 | 0.224 | -0.14, 0.59 | .225 | .507 |
| Corona radiata | Cho | 45 | 0.258 | -0.01, 0.53 | .063 | .284 |

**Supplementary Table 8.**
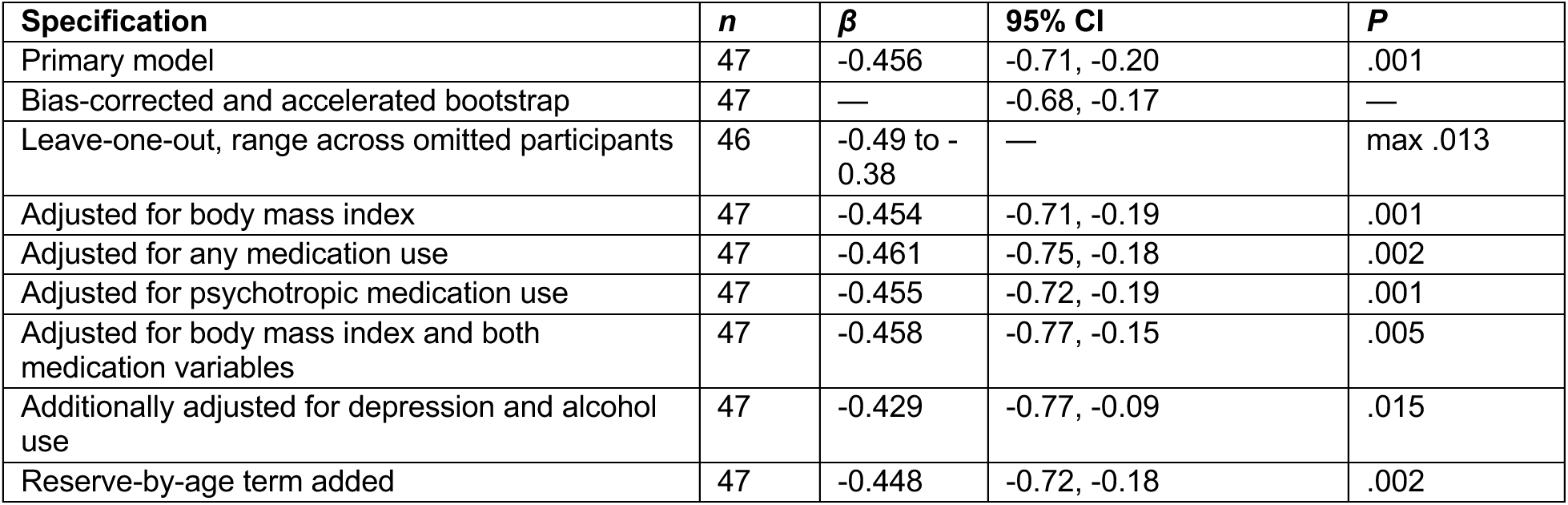
Sensitivity analyses for the reserve-by-PCG tNAA/tCr interaction on delayed recall. To determine whether the neurochemical interaction depends on distributional assumptions, individual participants or the covariates included, we re-estimated it under each alternative specification. All models adjust for injury burden, age, education and post-traumatic stress symptoms. The bias-corrected and accelerated bootstrap interval (10,000 resamples) excluded zero, with 99.8% of resampled estimates below zero. Leave-one-out estimates refit the model with each participant omitted in turn. In leave-one-out cross-validation, adding the interaction raised the cross-validated R² for delayed recall from 0.09 to 0.33.

| **Specification** | ***n*** | ***β*** | **95% CI** | ***P*** |
|---|---|---|---|---|
| Primary model | 47 | -0.456 | -0.71, -0.20 | .001 |
| Bias-corrected and accelerated bootstrap | 47 | — | -0.68, -0.17 | — |
| Leave-one-out, range across omitted participants | 46 | -0.49 to -0.38 | — | max .013 |
| Adjusted for body mass index | 47 | -0.454 | -0.71, -0.19 | .001 |
| Adjusted for any medication use | 47 | -0.461 | -0.75, -0.18 | .002 |
| Adjusted for psychotropic medication use | 47 | -0.455 | -0.72, -0.19 | .001 |
| Adjusted for body mass index and both medication variables | 47 | -0.458 | -0.77, -0.15 | .005 |
| Additionally adjusted for depression and alcohol use | 47 | -0.429 | -0.77, -0.09 | .015 |
| Reserve-by-age term added | 47 | -0.448 | -0.72, -0.18 | .002 |

**Supplementary Table 9.** Mapping of computerized subtests to the scored cognitive domains. This is the standard mapping supplied by the instrument. The fifteen domains analyzed, including Social Acuity from the Perception of Emotions Test, are those listed in Supplementary Table 6. Several domains derive from shared subtests and are therefore not statistically independent.

| **Cognitive test** | **Derived CNSVS domains** |
|---|---|
| Verbal Memory Test (VBM) | Composite Memory; Verbal Memory |
| Visual Memory Test (VSM) | Composite Memory; Visual Memory |
| Finger Tapping Test (FTT) | Psychomotor Speed; Motor Speed |
| Symbol Digit Coding (SDC) | Processing Speed; Psychomotor Speed |
| Stroop Test (ST) | Reaction Time; Complex Attention |
| Shifting Attention Test (SAT) | Cognitive Flexibility; Executive Function; Complex Attention |
| Continuous Performance Test (CPT) | Simple Attention; Complex Attention |
| Perception of Emotions Test (POET) | Social Acuity |
| Non-Verbal Reasoning Test (NVRT) | Reasoning |
| Four-Part Continuous Performance Test (FPCPT) | Working Memory; Sustained Attention |

**Supplementary Table 10.** Descriptions of the computerized subtests. Descriptions are provided verbatim from the CNS Vital Signs score report.

| **Cognitive test** | **Description** |
|---|---|
| Verbal Memory Test (VBM) | Subjects have to remember 15 words and recognize them in a field of 15 distractors. The test is repeated at the end of the battery. The VBM test measures how well a subject can recognize, remember, and retrieve words. Correct hits refers to the number of target words recognized. Low scores indicate verbal memory impairment. |
| Visual Memory Test (VSM) | Subjects have to remember 15 geometric figures and recognize them in a field of 15 distractors. The test is repeated at the end of the battery. The VSM test measures how well a subject can recognize, remember, and retrieve geometric figures. Correct hits refers to the number of target figures recognized. Low scores indicate visual memory impairment. |
| Finger Tapping Test (FTT) | A test of motor speed and fine motor control. There are three rounds of tapping with each hand. The FTT measures the speed and number of finger taps with each hand. Low scores indicate motor slowing. Speed of manual motor activity varies with handedness. |
| Symbol Digit Coding (SDC) | The SDC measures speed of processing and draws on several cognitive processes simultaneously, including visual scanning, visual perception, visual memory, and motor function. Errors may reflect impulsive responding, misperception, or confusion. |
| Stroop Test (ST) | The ST measures simple and complex reaction time, inhibition and disinhibition, and mental flexibility or directed attention. It assesses how well a subject adapts to a rapidly changing and increasingly complex set of directions. Prolonged reaction times indicate cognitive slowing. |
| Shifting Attention Test (SAT) | The SAT measures executive function: how well a subject recognizes set shifting and abstraction, and manages multiple tasks simultaneously. Subjects adjust their responses to randomly changing rules. The best scores combine high correct responses, few errors, and short reaction time. |
| Continuous Performance Test (CPT) | The CPT measures sustained attention or vigilance and choice reaction time. Most normal subjects obtain near-perfect scores. A long response time may suggest cognitive slowing. More than two errors may be clinically significant; more than four indicate attentional dysfunction. |
| Perception of Emotions Test (POET) | The POET measures how well a subject can perceive and identify specific emotions, testing identification of the positive emotions happy and calm and the negative emotions angry and sad. Reaction times are much longer than in other tests, indicating the complexity of the processes governing emotional acuity. |
| Non-Verbal Reasoning Test (NVRT) | The NVRT measures how well a subject perceives and understands visual or abstract information and recognizes relationships between visual-abstract concepts. It comprises 15 progressively harder matrices, each presented for 14.5 seconds. |
| Four-Part Continuous Performance Test (FPCPT) | A four-part test of working memory and sustained attention. Part one is a simple reaction time test. Part two is a variant of the continuous performance test requiring discrimination, generating choice reaction times. Part three is a one-back task. Part four is a two-back task and is used to measure working memory. Parts two to four are used to calculate the sustained attention domain. |

**Supplementary Table 11.**
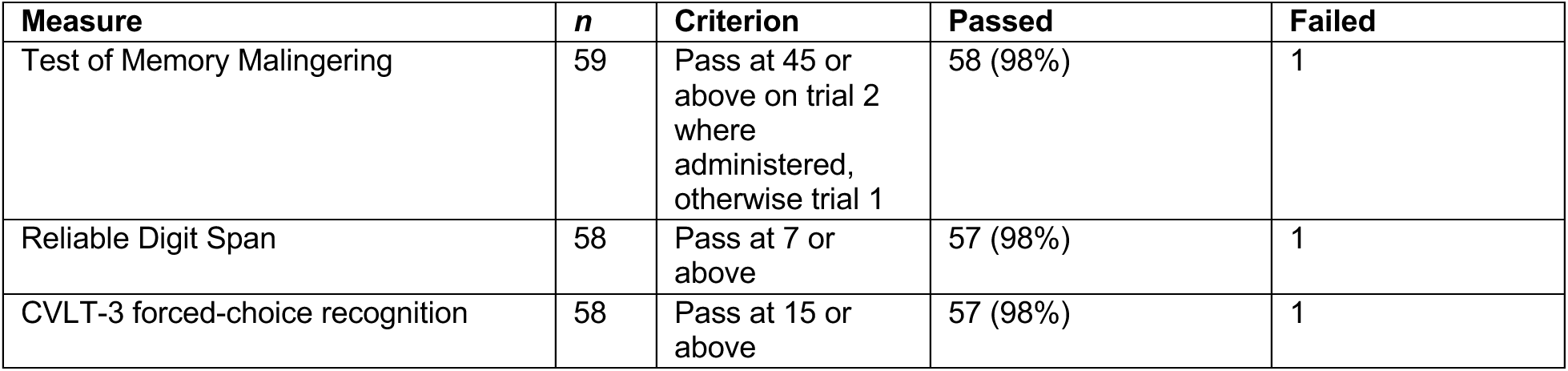
Performance validity. Injury exposure was self-reported, raising the possibility that variable engagement with testing contributed to the results. To address this, performance validity was assessed with three measures against the cutoffs shown. The reliable digit span row applies a score of 7 or above as a pass, following the more conservative of the two cutoffs evaluated by Schroeder et al. [67]. The Test of Memory Malingering row scores trial 2 where it was administered and trial 1 otherwise, since trial 2 is given only when trial 1 falls below criterion; three participants have no recorded administration and are not represented. The single failure is a participant whose trials were both discontinued because of an administrative error, giving scores of 11 and 17; errors on the first ten items alone predict performance validity [86], so the abbreviated administration is scored as a failure rather than treated as missing, although item-level responses were not available to compute the formal ten-item score; she passed both other indicators and is one of the three participants examined in Supplementary Table 12, where omitting her leaves the behavioral and neurochemical estimates unchanged. A sensitivity analysis omitting the participants the assessor flagged for review is reported in Supplementary Table 12.

**Supplementary Table 12.** Sensitivity analyses excluding participants with potential testing-validity concerns. Three participants were flagged for review because of a noisy testing environment, below-cutoff forced-choice recognition, or premature discontinuation of an effort measure. Each had valid data on at least two of three indicators and was retained in the primary analyses. Excluding them did not materially change the behavioral or neurochemical estimates. Additionally excluding the participant who failed reliable digit span strengthened the reserve-by-injury interaction (*β* = 1.74, 95% CI [0.85, 2.63], *P* < .001) and preserved the neurochemical interaction (*β* = −0.41, 95% CI [−0.67, −0.15], *P* = .003).

| **Analysis** | ***n*** | ***β*** | **95% CI** | ***P*** |
|---|---|---|---|---|
| Reserve x ≥10 injuries, retained (primary analyses) | 58 | 1.363 | 0.23, 2.49 | .019 |
| Reserve slope in highest-burden band, retained (primary analyses) | 58 | 1.253 | 0.42, 2.09 | .004 |
| Reserve x PCG tNAA/tCr, retained (primary analyses) | 47 | -0.456 | -0.71, -0.20 | .001 |
| Reserve x ≥10 injuries, omitted | 55 | 1.555 | 0.48, 2.62 | .005 |
| Reserve slope in highest-burden band, omitted | 55 | 1.446 | 0.65, 2.24 | <.001 |
| Reserve x PCG tNAA/tCr, omitted | 44 | -0.426 | -0.65, -0.20 | <.001 |

**Supplementary Table 13.**
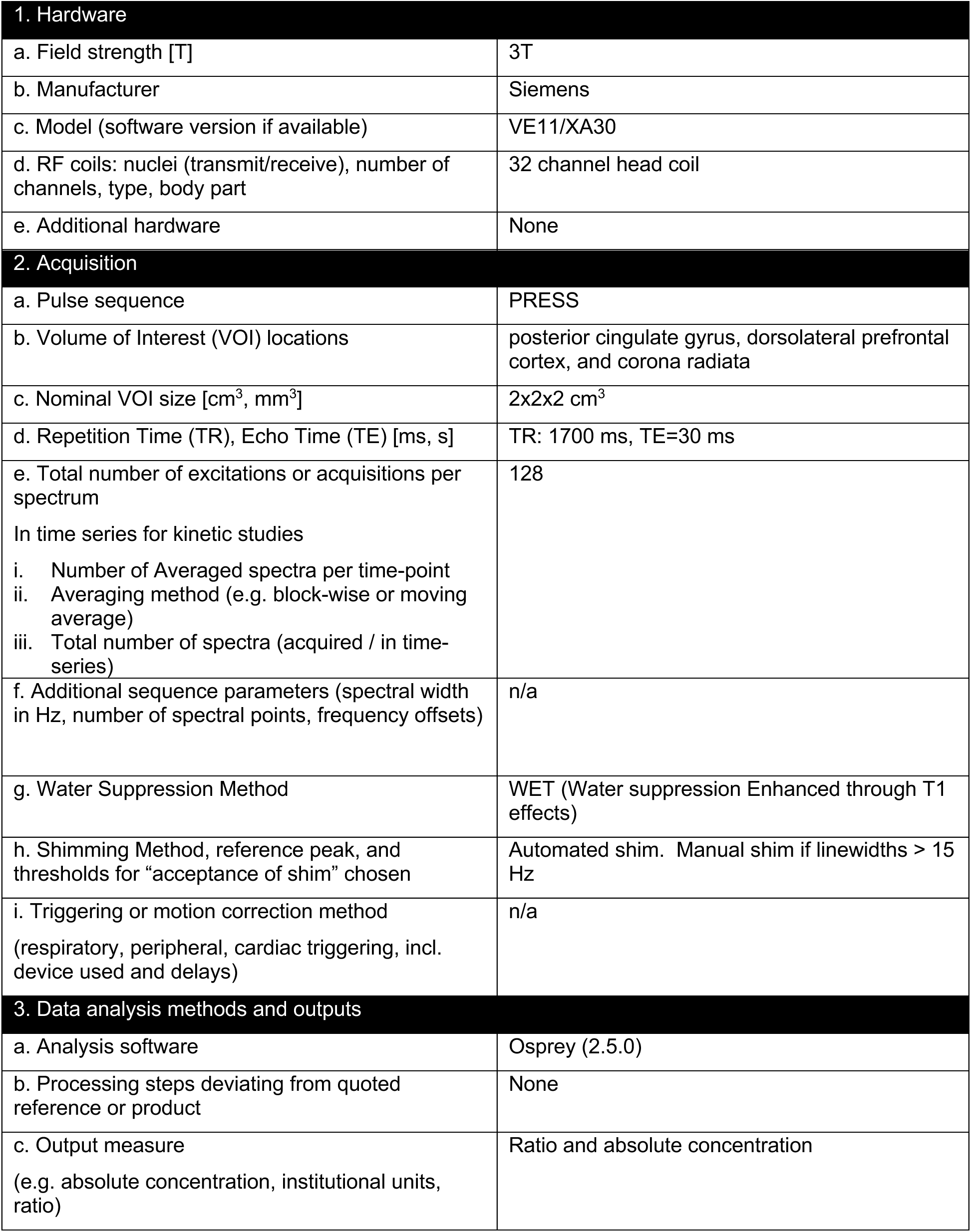
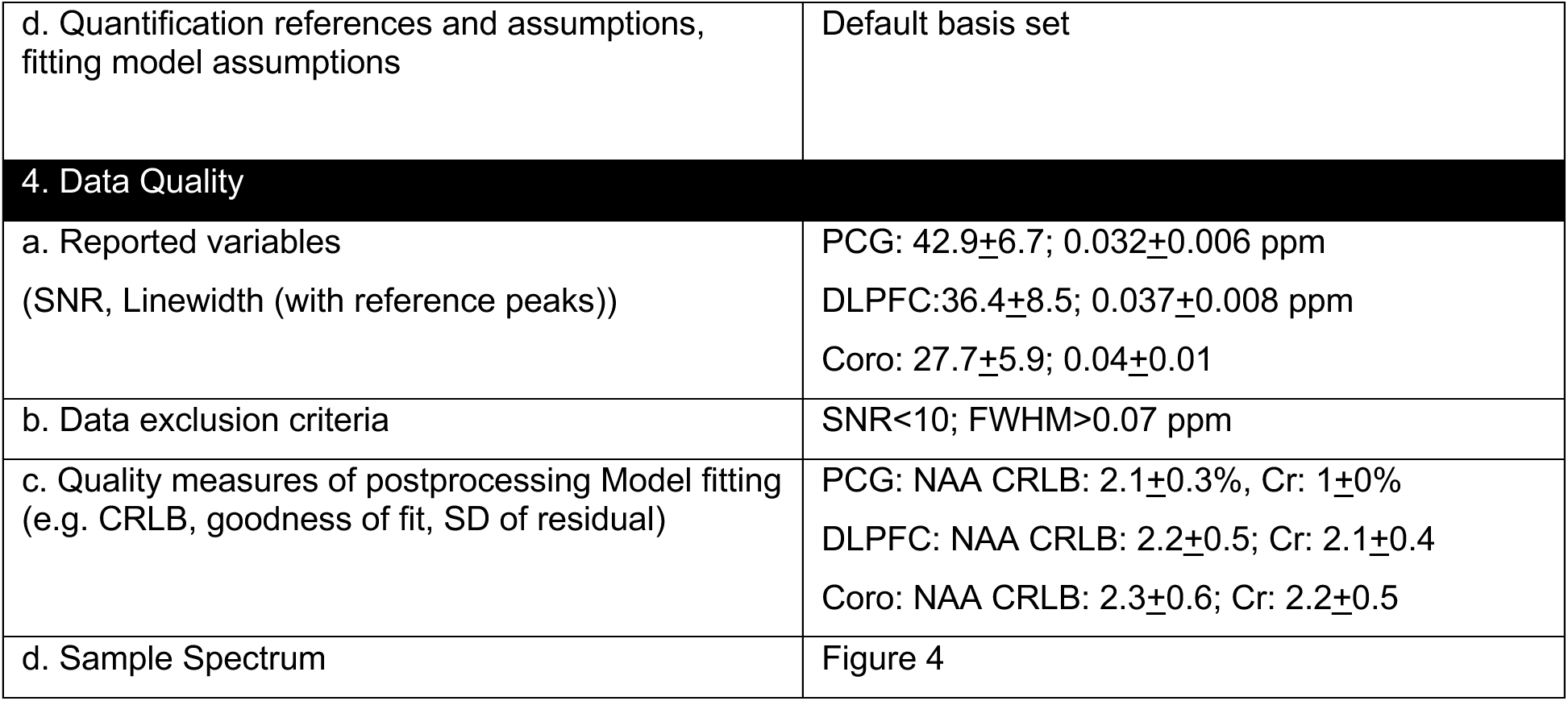
MRSinMRS checklist.

| 1. Hardware |  |
|---|---|
| a. Field strength [T] | 3T |
| b. Manufacturer | Siemens |
| c. Model (software version if available) | VE11/XA30 |
| d. RF coils: nuclei (transmit/receive), number of channels, type, body part | 32 channel head coil |
| e. Additional hardware | None |
| 2. Acquisition |  |
| a. Pulse sequence | PRESS |
| b. Volume of Interest (VOI) locations | posterior cingulate gyrus, dorsolateral prefrontal cortex, and corona radiata |
| c. Nominal VOI size [cm<sup>3</sup>, mm<sup>3</sup>] | 2x2x2 cm<sup>3</sup> |
| d. Repetition Time (TR), Echo Time (TE) [ms, s] | TR: 1700 ms, TE=30 ms |
| e. Total number of excitations or acquisitions per spectrum<br>In time series for kinetic studies<br>i. Number of Averaged spectra per time-point<br>ii. Averaging method (e.g. block-wise or moving average)<br>iii. Total number of spectra (acquired / in time-series) | 128 |
| f. Additional sequence parameters (spectral width in Hz, number of spectral points, frequency offsets) | n/a |
| g. Water Suppression Method | WET (Water suppression Enhanced through T1 effects) |
| h. Shimming Method, reference peak, and thresholds for “acceptance of shim” chosen | Automated shim. Manual shim if linewidths > 15 Hz |
| i. Triggering or motion correction method<br>(respiratory, peripheral, cardiac triggering, incl. device used and delays) | n/a |
| 3. Data analysis methods and outputs |  |
| a. Analysis software | Osprey (2.5.0) |
| b. Processing steps deviating from quoted reference or product | None |
| c. Output measure<br>(e.g. absolute concentration, institutional units, ratio) | Ratio and absolute concentration |
| d. Quantification references and assumptions, fitting model assumptions | Default basis set |
| 4. Data Quality |  |
| a. Reported variables<br>(SNR, Linewidth (with reference peaks)) | PCG: 42.9±6.7; 0.032±0.006 ppm<br>DLPFC:36.4±8.5; 0.037±0.008 ppm<br>Coro: 27.7±5.9; 0.04±0.01 |
| b. Data exclusion criteria | SNR<10; FWHM>0.07 ppm |
| c. Quality measures of postprocessing Model fitting (e.g. CRLB, goodness of fit, SD of residual) | PCG: NAA CRLB: 2.1±0.3%, Cr: 1±0%<br>DLPFC: NAA CRLB: 2.2±0.5; Cr: 2.1±0.4<br>Coro: NAA CRLB: 2.3±0.6; Cr: 2.2±0.5 |
| d. Sample Spectrum | Figure 4 |

## Notes

https://fitbir.nih.gov/

